# Localising enzymes to biomolecular condensates increases their accumulation and benefits engineered metabolic pathway performance in *Nicotiana benthamiana*

**DOI:** 10.1101/2025.01.29.635479

**Authors:** Anya L. Lindström Battle, Angus W. Barrett, Mark D. Fricker, Lee J. Sweetlove

## Abstract

Establishment of *N. benthamiana* as a robust biofactory is complicated by issues such as product toxicity and proteolytic degradation of target proteins / introduced enzymes. Here we investigate whether biomolecular condensates can be used to address these problems. We engineered biomolecular condensates in *N. benthamiana* leaves using transient expression of synthetic modular scaffolds. The *in vivo* properties of the condensates that resulted were consistent with them being liquid-like bodies with thermodynamic features typical of multicomponent phase-separating systems. We show that recruitment of enzymes to condensates *in vivo* led to several fold yield increases in one- and three-step metabolic pathways (citramalate biosynthesis and poly-3-hydroxybutyrate (PHB) biosynthesis, respectively). This enhanced yield could be for several reasons including improved enzyme kinetics, metabolite channelling or avoidance of cytotoxicity by retention of the pathway product within the condensate, which was demonstrated for PHB. However, we also observed a several-fold increase in amount of the enzymes that accumulated when they were targeted to the condensates. This suggests that the enzymes were more stable when localised to the condensate than when freely diffusing in the cytosol. We hypothesise that this stability is likely the main driver for increased pathway product production. Our findings provide a foundation for leveraging biomolecular condensates in plant metabolic engineering and advance *N. benthamiana* as a versatile biofactory for industrial applications.

## 1. Introduction

*Nicotiana benthamiana* has been used as a model to study fundamental questions in plant biology for several decades, especially by virtue of its use as a system for routine transient gene expression in leaf tissue. The ability to co-infiltrate more than 10 different *Agrobacterium* strains facilitates the reconstruction of complex metabolic pathways [1]. The rapid and high-level expression levels achievable through agroinfiltration has led to the emergence of *N. benthamiana* as a bioproduction species for both high-value proteins and chemicals, with research ongoing to further optimise the agroinfiltration process including by using whole-plant vacuum infiltration [2] and self-replicating viral vectors [3].

Despite the power of transient expression in *N. benthamiana* certain challenges remain in exploiting *N. benthamiana* as a biofactory. One of these is the toxicity associated with high accumulation levels of some products. In nature, plants can accumulate some natural products to remarkably high levels; for example, young sorghum seedlings accumulate the cyanogenic glucoside dhurrin to 30 % of their dry mass [4], vanillin glucoside is present at a concentration of 4.7 M in *Vanilla planifolia* pods [5], and *Cannabis sativa* accumulates cannabinoids to concentrations approaching 0.3 M [6]. Toxicity is avoided in these examples by compartmentation of the product, for instance in the tip of trichomes in cannabis or in senescent chloroplasts in vanilla. Compartmentation of compounds is also important to prevent modification by endogenous enzymes such as hydrolases, oxidoreductases, or transferases which form part of endogenous *N. benthamiana* metabolism or its xenobiotic detoxification system [7]. In fact the production and storage of toxic or labile compounds is a key concern in the metabolic engineering of all hosts [8].

In addition to classical membrane-bound organelles, it is now well-established that cells also organise and concentrate components into nonmembrane-bound structures such as nucleoli, Cajal bodies, and pyrenoids [9]. These bodies are collectively termed biomolecular condensates because they spatially concentrate biomolecules [10], and have a range of underlying biophysical properties from phase-separated liquid-liquid droplets through to gel-like structures and even to crystalline entities. These properties are condition-dependent and can change over time [11]. In addition to the well-documented ability of such structures to concentrate proteins [12], recent work has shown they also sequester small molecules including ATP and other metabolites [13–15]. In fact, it has been suggested that many plant bioactive compounds are stored in biomolecular condensates composed of natural deep eutectic solvents, including cannabinoids and dhurrin [16], and that enzymes remain functional inside these structures [17].

*In vitro* studies have also shown that enzymes localised to biomolecular condensates can gain kinetic benefits [18]. This effect may arise from the recruitment of enzymes into substructures with varying densities within the condensates, which could promote favourable orientations of enzymes and substrates [19].

The ability of biomolecular condensates to selectively concentrate proteins and metabolites and favourably influence enzyme kinetics has resulted in the design of synthetic condensates as a compartmentation tool for metabolic engineering. This commonly involves re-purposing proteins shown to phase-separate *in vitro*, such as those with intrinsically disordered regions (IDRs). IDRs usually possess multiple repeated domains containing residues able to form the many reversible interactions required for condensate formation [20]. For example, the RGG IDR from the *Caenorhabditis elegans* LAF-1 protein has been shown to stably phase-separate *in vitro* [21] and its role in driving phase-separation during P-granule formation is well-documented [22]. This domain has therefore been used to generate several synthetic RGG-based biomolecular condensates in yeast and human cells, with favourable outcomes for metabolic engineering [23–26]. However, although synthetic condensates have recently been reported in plant cells [27], the potential of these compartments for plant metabolic engineering has not yet been explored.

Another challenge for heterologous protein production in *N. benthamiana* is proteolysis of the target protein / enzyme, to the extent that proteolytic degradation is often cited as a bottleneck to the industrial viability of agroinfiltration [28–30]. Plant genomes encode several hundred proteolytic enzymes most, but not all, of which reside in the apoplast or the vacuole [31], and the expression of protease-encoding genes has been shown to significantly increase upon agroinfiltration as a result of a pathogen defence response [28]. Target proteins such as IgG antibodies, cytokines or plasminogen activators are often reported to be affected by proteolytic degradation when produced in plant cells [32–34]. It is likely that proteolysis also affects enzymes overexpressed for product accumulation in metabolic engineering attempts, thus negatively affecting the accumulation of a desired end product.

Attempts at overcoming proteolytic degradation have included the treatment of plants with chemical protease inhibitors [35, 36], the co-expression of target proteins with protease inhibitors [37, 38], or the knockdown of protease-encoding genes [39]. However, it is often challenging to identify exactly which protease is responsible for the observed degradation and hence how to inhibit it [29, 30]. One of the most promising approaches for protecting heterologous proteins from degradation is the development of targeting strategies to sequester target proteins away from hydrolytic cellular compartments. For instance, fusing target proteins to the N-terminal domain of the maize seed storage protein γ-zein (Zera) led to their accumulation in ER-derived protein bodies, increasing the accumulation of the Zera-fused subunit vaccine F1-V from *Yersinia pestis* by 3-fold in transiently transformed *N. benthamiana* [40]. Similarly targeting the transiently expressed 14D9 antibody to the vacuole increased yield 10- to 15-fold [41]. Proteins have also been targeted to the plastid and to the apoplast [42]. However, these compartmentation strategies are not ideal for metabolic engineering as they involve subcellular locations that are not favourable for metabolism.

In this paper we test the hypothesis that targeting enzymes to biomolecular condensates could help address the needs for compartmentation and proteolysis-avoidance for metabolic engineering in *N. benthamiana*. We build synthetic biomolecular condensates in *N. benthamiana,* interrogate their biophysical properties, and show that the localisation of heterologous enzymes to biomolecular condensates increases product yield from both a one- and a three-step metabolic pathway. The localisation of these enzymes to biomolecular condensates dramatically increased their abundance compared to conventional over-expression and we suggest that this is at the root of the observed increased product yield. Finally, we present evidence that desirable end products can be enriched inside these biomolecular condensates. The findings presented here have promising implications for the ongoing development of *N. benthamiana* as a biofactory for valuable proteins and compounds.

## 2. Results and Discussion

### 2.1. Building synthetic biomolecular condensates in N. benthamiana

A construct for the creation of synthetic biomolecular condensates in *N. benthamiana* was designed based on previous work in *Saccharomyces cerevisiae* [26]. The system is composed of a scaffold and a client; the design of the client will be covered in more detail in section 2.3. The scaffold was built from two copies of RGG, the IDR from the *C. elegans* LAF-1 protein. The monomeric green/yellow GFP-derived fluorescent protein mClover3 was included for visualization [43], and one of a cognate pair of synthetic coiled-coil peptides (SYNZIP1) was included for client recruitment [44]. Monomeric fluorescent proteins are recommended in this type of experiment as multimerising activity might affect condensate formation [45]. For purification purposes, both the scaffold and client included a 10×His tag on the C or N terminus, respectively. The client and scaffold system was encoded as a polycistronic construct with the protein units separated by the InteinF2A self-cleaving fusion protein domain, composed of a *Ssp* DnaE mini-intein variant engineered for hyper-N-terminal autocleavage which was covalently linked to the 2A peptide from the foot-and-mouth disease virus [46]. After translation, the scaffold and client should form two separate fusion proteins; the scaffold protein should assemble into biomolecular condensates via the multiple and weak interactions between RGG domains and is illustrated in Figure 1A.

**Figure 1:**
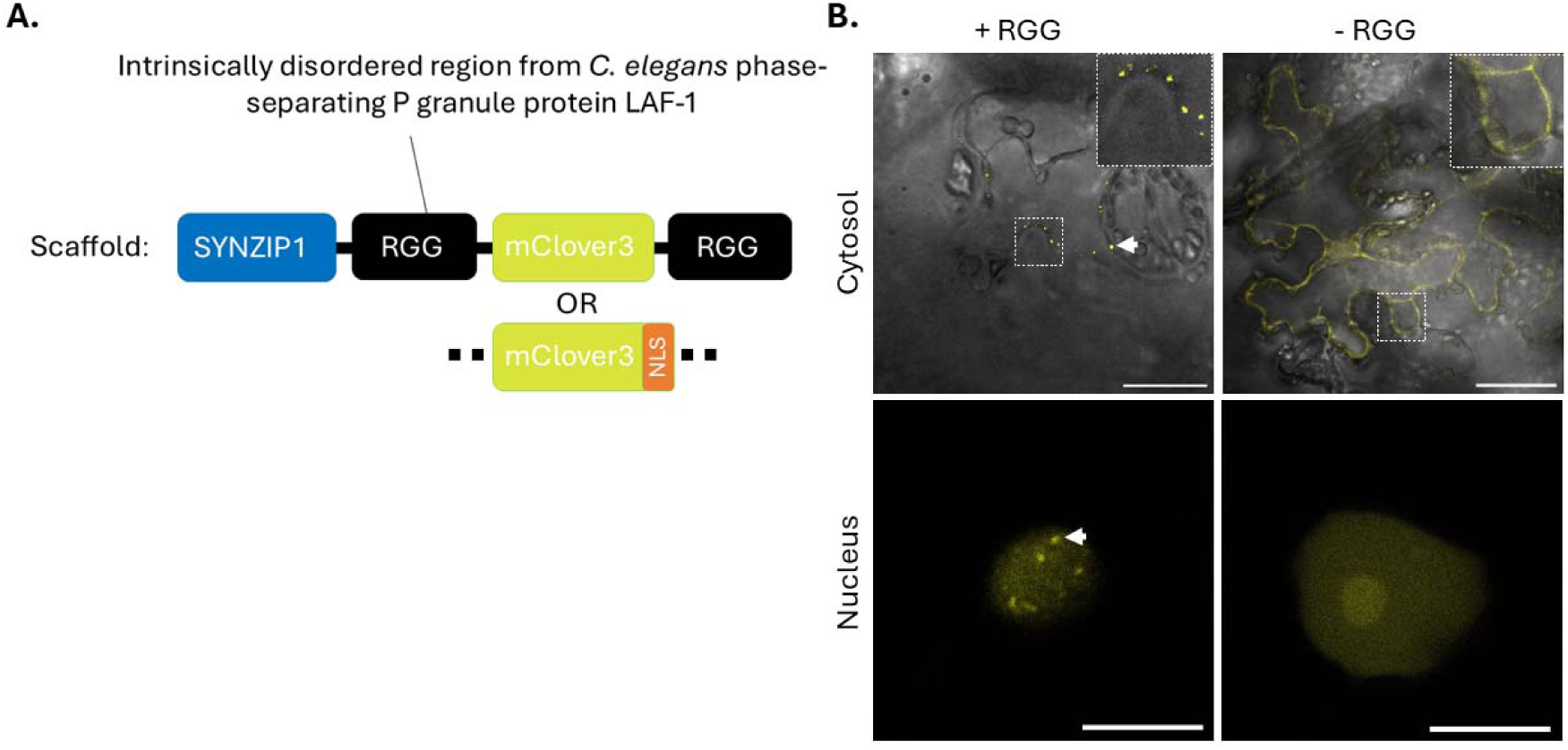
Building synthetic biomolecular condensates in N. benthamiana. (A) the design of a scaffold construct for building synthetic biomolecular condensates in plants. (B) The RGG-dependent construction of synthetic biomolecular condensates in the nucleus and in the cytosol of N. benthamiana. Biomolecular condensates indicated by white arrows. Scale bar is 50 μm (cytosol) and 10 μm (nucleus).

Two versions of the scaffold were designed, the difference being the presence or absence of a nuclear localisation signal (NLS) on the C-terminus of mClover3. In addition, both versions of the construct were also expressed without the RGG domains as a control (not expected to form condensates). Constructs were carried within the pK7WG2 Gateway plant expression vector which places the construct downstream of the constitutive p35S promoter [47].

The nuclear and cytosolic versions of the construct were transformed into mature *N. benthamiana* leaves and expression of mClover3 was monitored over 7 d using confocal laser scanning microscopy. Punctate structures that were putative biomolecular condensates were observed in both nuclei and cytosol of transiently transformed *N. benthamiana* leaves when infiltrated with the nuclear or cytosolic version of the construct, respectively (Figure 1B). Nuclear localisation was confirmed using DAPI staining (Figure S1A). Biomolecular condensates were first visible 2 days post-infiltration (dpi). In both subcellular locations the mClover3 signal increased until 3 or 5 dpi and then decreased, such that barely any signal was visible after day 7 (Figure S1B). The exception to this was in some cells transformed with the cytosolic version of the construct where a small number of very large biomolecular condensates were occasionally still seen 7 dpi. All subsequent characterisation was carried out on leaf tissue at 5 dpi unless otherwise specified.

To confirm that the punctate structures were forming due to interactions between RGG domains, versions of the nuclear and cytosolic constructs without these domains (referred to as – RGG) were infiltrated into *N. benthamiana* leaves, and expression over 7 d was monitored as described. In both the cytosol and the nucleus diffuse mClover3 signal was observed which increased until 3 dpi (Figure S1B). No punctate structures were observed in plants infiltrated with these constructs. A single large and diffuse structure was observed in some nuclei, posited to be the nucleolus (Figure 1B).

The abilities of the cytosolic and nuclear versions of the construct to form biomolecular condensates were compared by calculating the number and volume of biomolecular condensates in each cell (Figure S1C). There was a large difference (*P* < 0.001) between the two locations for both metrics; on average, there were 7.7 biomolecular condensates per nucleus, but 41.9 in the cytosol. In some cases, in the cytosol, there were up to 175 biomolecular condensates in a single cell. The volume of the biomolecular condensates was also significantly larger in the cytosol than in the nucleus. The mean volume of a nuclear biomolecular condensate was 0.47 μm^3^ whereas the mean cytosolic biomolecular condensate volume was 5.7 μm^3^. The largest cytosolic biomolecular condensate measured was 495 μm^3^. In plant cells the volume of the cytosol is expected to be approximately 30-fold larger than the volume of the nucleus [48]; as such it is not unexpected that it can accommodate both more and larger biomolecular condensates. In addition, for biomolecular condensates to be formed in the nucleus the scaffold protein must be trafficked into the nucleus which introduces an extra obstacle to biomolecular condensate formation.

### 2.2. A pipeline for the in vivo characterisation of synthetic biomolecular condensates

Having built synthetic biomolecular condensates in *N. benthamiana*, we wanted to explore their potential usefulness as a tool for metabolic engineering. This necessarily involves an understanding of their biophysical properties. Although no comprehensive framework exists with which to predict whether or how a given substance will diffuse into and out of a given condensate, whether the structure is solid- or liquid-like will affect its permeability by changing the energetic barrier to entering or leaving the biomolecular condensate for substrates, cofactors, or enzymes [49]. This in turn might affect the activity of any recruited enzyme and hence the suitability of a synthetic condensate for metabolic engineering. Here we have developed an easy-to-use pipeline for the analysis of synthetic biomolecular condensates in *N. benthamiana* leaves which could help inform future attempts at building synthetic condensates in this species. The code for running the analysis is available at https://github.com/AnyaLindstromBattle/Biomolecular_condensates_N_benthamiana_PBJ2025.

If a condensate is liquid-like, it likely forms through a phenomenon known as liquid-liquid phase separation (LLPS) [50]. In its simplest form this involves a solute demixing into two stable liquid phases, one solute-rich and one solute-poor, because of transient, low-affinity interactions according to a set of well-defined thermodynamic rules [45]. However, more recently it has become clear that not all condensates identified in living systems are liquids and can form through mechanisms distinct from LLPS [11].

It is very difficult to demonstrate LLPS *in vivo*, and caution should be exercised in assuming that condensates are being formed through LLPS without the required due diligence needed to establish whether phase-separation is actually occurring [11, 51]. *In vitro* there exist rigorous standards by which one may determine whether a system is undergoing LLPS, but the intracellular environment is much more complex than any *in vitro* system and involves the coexistence of many thousands of small molecules with potential for homo- or heterotypic interactions in a small volume. Recent work has sought to identify a set of minimum experiments to support identification of LLPS *in vivo.* These include evidence for the fusion of spherical structures, demonstration of a concentration dependence of condensation, and a test of whether LLPS-deficient mutants identified *in vitro* assemble in cells [52].

We sought to characterise the nuclear RGG-based synthetic biomolecular condensates built in this work according to these three metrics. Although spherical nuclear biomolecular condensates were observed (Figure 2A) we were unable to observe any fusion events. These events may be rare and may require long observation times. However we were able to demonstrate that LLPS-deficient mutants identified *in vitro* behaved as expected, with the no-RGG versions of the scaffold unable to form biomolecular condensates (Figure 1B). Note that although fluorescence recovery after photobleaching (FRAP) is a contentious way of supporting LLPS [11, 53], we also showed recovery of fluorescence in both nuclear and cytosolic condensates supporting at least some mobility of the scaffold molecule (Figure S1C).

**Figure 2:**
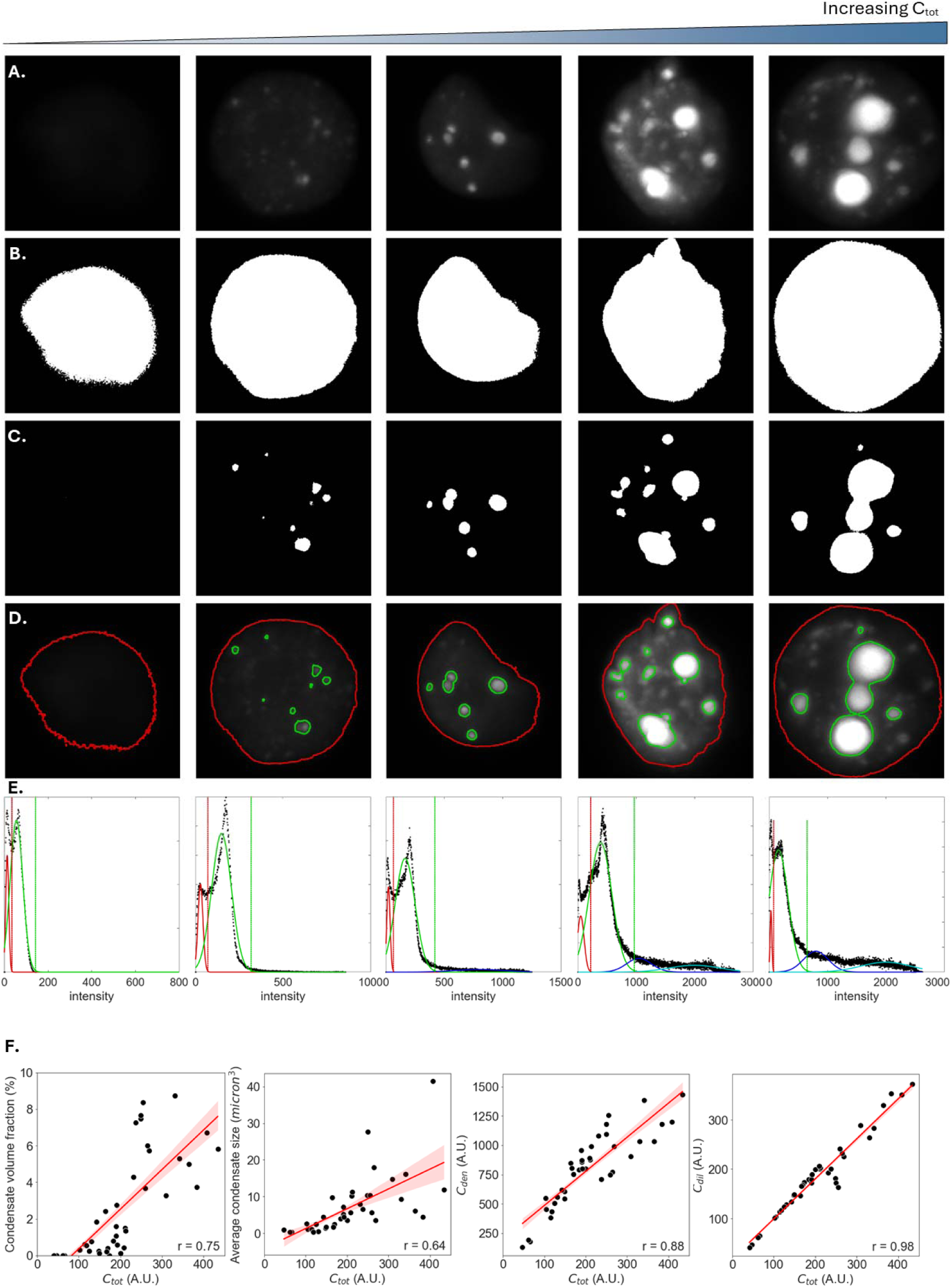
The in vivo characterisation of synthetic biomolecular condensates targeted to the nucleus. (A) Transient expression in N. benthamiana generated nuclei with a range of total nuclear scaffold concentrations (C_tot_) . Images were segmented into (B) the nucleus and (C) the condensates, with (D) showing the outlines of each segmented region on the original image. (E) Segmentation was achieved by fitting Gaussian mixed models to signal intensity distributions where the red curve fits the background, the green the nucleoplasm, and the blue and cyan (depending on how many Gaussians provides the best fit of the data, see Experimental procedures) the biomolecular condensate signal. The vertical lines show the value at mean + 3×standard deviations (SD) of the different Gaussians and was used to determine the intensity threshold for signal pertaining to the nucleus (red, mean+3×SD from the Gaussian fitted to the background) or to the condensates (green, mean+3×SD from the Gaussian fitted to the nucleoplasm). Note that for clarity these images are maximum projections but the analysis was carried out on full z-stack images. (F) The relationship between C_tot_ and condensate volume fraction, average condensate size, the concentration of scaffold in the dense phase (i.e. the condensates, C_den_), and the concentration of scaffold in the dilute phase (i.e. the nucleoplasm, C_dil_). Shaded area represents SEM, and the Pearsons correlation coefficient (r) is shown in the bottom-right corner.

Finally, we used microscopy-based techniques to interrogate whether the RGG-based nuclear biomolecular condensates showed a concentration-dependence of condensation. This took advantage of the heterogeneous gene expression levels expected across an agroinfiltrated *N. benthamiana* leaf and was based on recent work in mammalian, bacterial, and viral cells to test concentration-dependent thermodynamic effects and support the identification of LLPS *in vivo* [54–56]. Randomly selected nuclei were imaged 3 and 4 dpi with identical acquisition settings, resulting in 42 nuclear images with a range of condensate-forming behaviours (Figure 2A). Images were segmented using Gaussian mixed models (GMMs) (Figures 2B-E) and the total nuclear concentration of the scaffold (C_tot_), the concentration of the scaffold in the nucleoplasm (i.e. the dilute phase, C_dil_), and the concentration of the scaffold in the condensates (i.e. the dense phase, C_den_) were calculated for each image using the fluorescence level of mClover3. In addition, we calculated the condensate volume fraction and the average condensate volume in each image (Figure 2F). This allowed for an exploration of concentration-dependent effects on condensate formation.

We showed that both condensate volume fraction and average condensate volume increased with C_tot_, as would be expected for condensates formed through LLPS [52]. However, in simple binary phase separation the C_den_ and C_dil_ should reach a constant level, referred to as the saturation concentration (C_sat_), even under varied protein expression levels once LLPS has occurred. However, neither the concentration of the dilute or the dense phases remained fixed but instead increased with an increase in C_tot_ (Figure 2F).

These data are incompatible with the behaviour expected of a simple binary phase separating system. Instead, they are consistent with the more richly textured thermodynamics of multicomponent phase-separating systems as identified in other biomolecular condensates *in vivo,* such as the nucleolus [54]. In multicomponent systems, the phase-separating behaviour is not only governed by the homotypic interactions between the scaffold molecules, but is also affected by heterotypic interactions between the scaffold and other element(s) [54]. The polycistronic scaffold-client construct used in this study involves the simultaneous expression of both a scaffold molecule and a client molecule (in this case, mCherry, as detailed in section 2.3). Although the client was not directly imaged in this analysis, it is likely being recruited to the nuclear biomolecular condensates. Thus, in addition to scaffold-scaffold interactions, it is reasonable to assume that client-scaffold interactions also influence the phase-separating behaviour of the system.

### 2.3. Functionalisation of synthetic biomolecular condensates for metabolic engineering

To determine whether synthetic biomolecular condensates could be useful compartmentation tools for metabolic engineering in *N. benthamiana* we tested whether proteins could be targeted to them and retain catalytic activity. The scaffold-client system as described in section 2.1 results in client recruitment to the scaffold-based biomolecular condensate via the dimerisation of a synthetic coiled-coil pair (SYNZIP1 [fused to the scaffold] and SYNZIP2 [fused to the client] [44]). Three different client designs were tested (Figure 3A). In the first, the client comprised the monomeric red fluorescent protein mCherry [57] fused to SYNZIP2. There was substantial overlap of fluorescence signals from mClover3 and mCherry in both cytosol- and nucleus-targeting constructs indicating significant co-localisation of these proteins (Figure 3B). This was statistically verified using a Pearson correlation analysis (Figure S1E). That mCherry signal was detected in the nucleus suggests that client recruitment via SYNZIP dimerisation may be occurring in the cytosol before recognition of the NLS on the scaffold and subsequent trafficking into the nucleus (since the mCherry-SYNZIP2 protein lacked a NLS). As proteins of the correct size for the client and scaffold part of the construct were detected in immunoblots against the leaf crude lysate (Figure S1F), we concluded that the polycistronic construct was cleaving as expected, and that the interaction between SYNZIP1 and SYNZIP2 was effectively recruiting the mCherry client into the structure formed by the scaffold.

**Figure 3:**
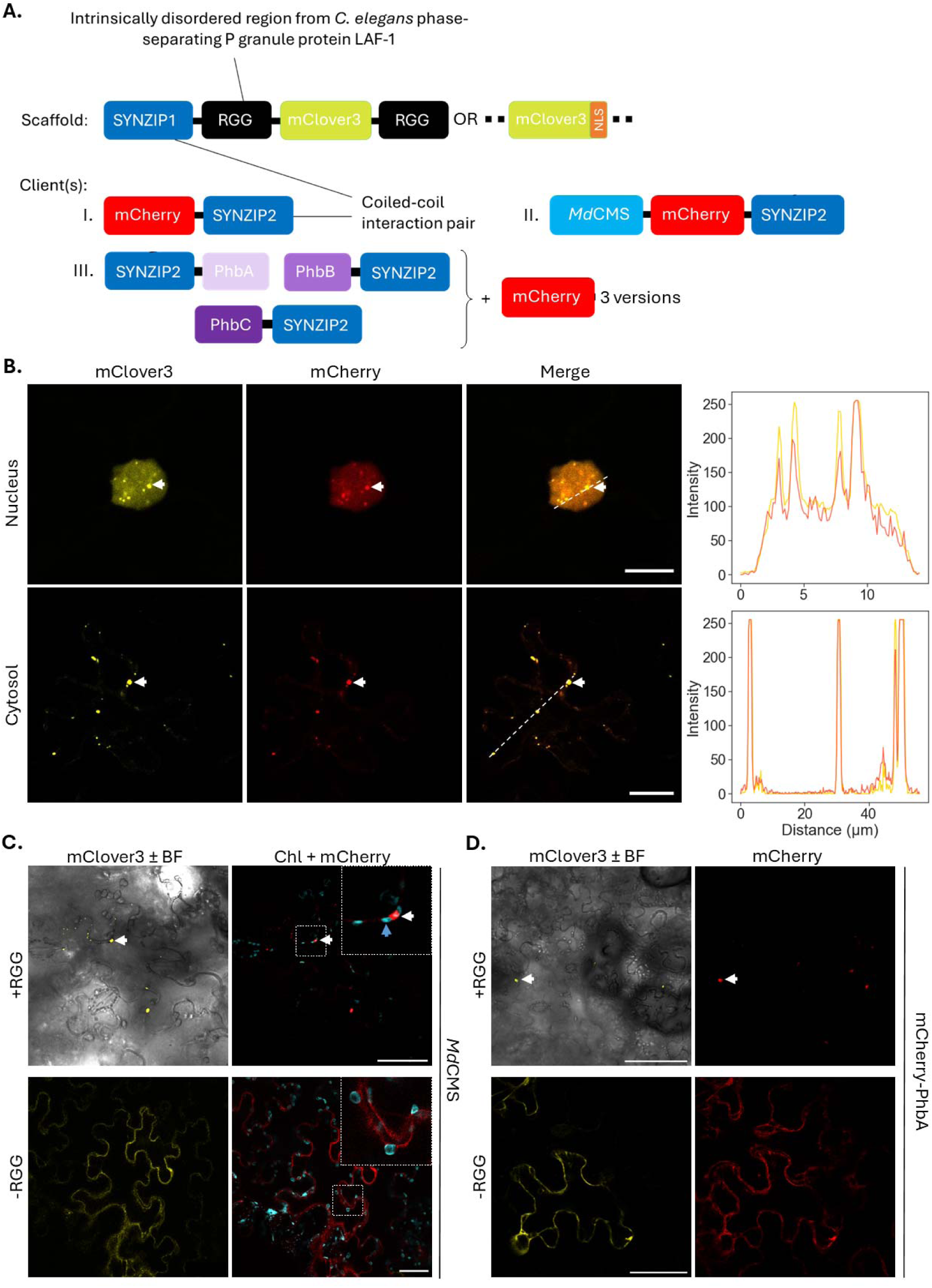
Functionalisation of synthetic biomolecular condensates for metabolic engineering. (A) An illustration of the scaffold-client system designed and used to generate synthetic biomolecular condensates in plants to which mCherry (I.), MdCMS (II.), or the PHB biosynthetic pathway (III.) is targeted. (B) Left: Punctate fluorescence indicates likely biomolecular condensate formation and mCherry recruitment in both nuclei and cytosol of agroinfiltrated N. benthamiana leaves, as indicated using a white arrow. Right: Signal intensity of both mClover3 (yellow) and mCherry (red) plotted across a transect indicated as a dashed white line in the merged image. Scale bar is 10 μm (nucleus) and 20 μm (cytosol). (C) Co-localisation of mClover3 and mCherry-fused cytosolic MdCMS. White arrow indicates a condensate, and blue arrow indicates a chloroplast. The removal of the RGG domain ablates biomolecular condensate formation and results in diffuse mClover3 and mCherry cytosolic signal. Scale bar is 100 μm. (D) Co-localisation of mClover3 and mCherry-fused β-ketothiolase (PhbA). White arrow indicates a condensate. The removal of the RGG domain ablates biomolecular condensate formation and results in diffuse mClover3 and mCherry cytosolic signal. For PhbB and PhbC please see Figure S3. Scale bar is 100 μm.

The second client tested comprised the enzyme citramalate synthase from *Malus domestica* (*Md*CMS). Targeting only to cytosolic condensates was examined. *Md*CMS catalyses the condensation of acetyl-CoA and pyruvate to citramalate [58] (Figure 4A); citramalate is not expected to be present in WT *N. benthamiana* facilitating its quantification in agroinfiltrated tissue. The native *Md*CMS is localised to the plastid [58]; to construct a version of *Md*CMS that would remain in the cytosol to be targeted to the condensates, the N-terminal chloroplast transit peptide was identified and removed (Figure S2).

**Figure 4:**
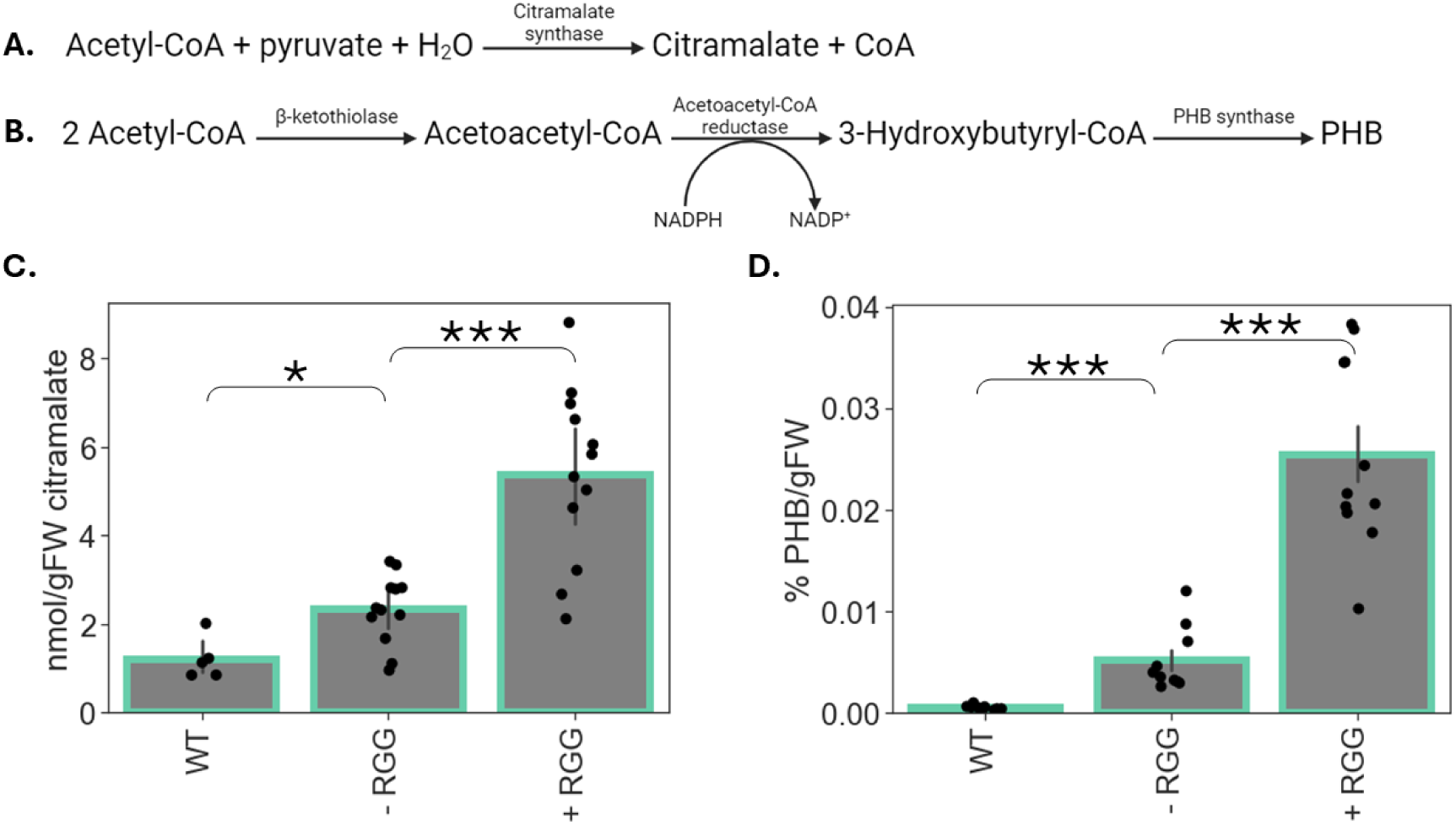
Localising enzymes to biomolecular condensates increases product yield. (A) The pathway for the production of citramalate by citramalate synthase. (B) The pathway for the production of PHB by the three-enzyme PHB pathway from R. eutropha. (C) Citramalate contents in un-infiltrated leaves (WT) or in leaves 5 dpi with either the RGG-containing (+ RGG) or no-RGG-containing (- RGG) scaffold construct. n = 5-12. (D) PHB contents in un-infiltrated leaves (WT) or in leaves 13 dpi with either the RGG- containing (+ RGG) or no-RGG-containing (- RGG) scaffold construct. n = 10-11. For both (E) and (F) error bars represent SEM; ‘***’ represents a P-value < 0.001, ‘**’ a P-value < 0.01, ‘*’ a P- value < 0.05, or ‘n.s.’ a P-value > 0.05 (Mann-Whitney U).

The scaffold-client system encoding *Md*CMS without the putative chloroplast transit peptide (Figure 3A) was transiently transformed into *N. benthamiana* leaves and imaged 5 dpi. Cytosolic biomolecular condensates were visible and mCherry was co-localised with mClover3 signal indicating that the mCherry-*Md*CMS fusion was being recruited to the synthetic condensates (Figure 3C). No mCherry signal was observed in the chloroplasts showing the predicted sequence to be a true chloroplast targeting signal. Removing the RGG domains resulted in no biomolecular condensate formation and the mClover3 and mCherry signals were instead found to be diffusely present in the cytosol (Figure 3C).

Finally, we wanted to test whether more than one enzyme could be targeted to cytosolic biomolecular condensates and for this we selected the three enzymes of the poly-3- hydroxybutyrate (PHB) pathway, a biodegradable polymer synthesized by various eubacteria as a carbon storage compound [59]. In *Ralstonia eutropha*, the PHB biosynthesis pathway involves β-ketothiolase, acetoacetyl-CoA reductase, and PHB synthase to convert acetyl-CoA to PHB [60] (Figure 4B). Rather than expressing the scaffold and client as part of the same polycistronic construct, we took advantage of the ability for combinatorial co-infiltration of *Agrobacterium* strains carrying different constructs to design a system where the scaffold and clients were encoded within separate plasmids. The scaffold was designed as previously described but was now individually expressed from a p35S promoter on pK7WG2. To generate the client construct, the three different PHB-producing enzymes were each fused to SYNZIP2 on either the N- (β-ketothiolase) or the C-terminus (acetoacetyl-CoA reductase and PHB synthase) and the three fusion proteins were separated by the self-cleaving viral peptide InteinF2A as previously described. The scaffold-client system modified for PHB production is illustrated in Figure 3A.

It was necessary to show that all three SYNZIP2-tagged PHB-producing enzymes could be recruited to the biomolecular condensate. To do this the client-containing construct was modified to create three versions where in each case a different enzyme was fused to mCherry. This allowed the ability of the tagged enzyme to be recruited to the biomolecular condensate in the presence of the other two enzymes to be tested whilst avoiding the need for simultaneous imaging of four different fluorophores. This method has been previously used in other systems to demonstrate the localisation of several different enzymes to condensates inside cells [61]. All three enzymes could be successfully recruited to the biomolecular condensate in the presence of the other two (Figure 3D and S3). In addition, and as previously shown, removing the RGG domains on the scaffold prevented biomolecular condensate formation and resulted in diffuse cytosolic signal from both mClover3 and mCherry (Figures 3D and S3).

These results showed that both fluorescent proteins and one or several enzymes could be successfully targeted to synthetic biomolecular condensates in *N. benthamiana*.

### 2.4. Targeting enzymes to biomolecular condensates increases product yield

For biomolecular condensates to be useful compartmentation tools in metabolic engineering enzymes need to remain functional when localised to them. The scaffold-client with *Md*CMS was hence used to determine whether enzyme activity was retained when targeted to cytosolic RGG-based biomolecular condensates. Samples were taken 5 dpi from *N. benthamiana* leaves infiltrated with the + RGG and – RGG versions of the *Md*CMS scaffold-client construct, and citramalate extracted and quantified (Figure 4C). Surprisingly, a small amount of citramalate was detected in WT tissue, even though *N. benthamiana* leaves are not known to produce citramalate. However, both versions of the construct resulted in higher (P < 0.05) levels of citramalate accumulation than WT showing that the enzyme was active both in the cytosol and when targeted to the condensate.

To determine whether the PHB-producing enzymes were functional when targeted to the condensates, the scaffold- and PHB client-containing *Agrobacterium* cultures were agroinfiltrated into *N. benthamiana* leaves and PHB was extracted and quantified. No PHB was detected in leaves 3 or 5 dpi. To determine whether quantifiable levels of PHB were achievable, the same leaves were re-infiltrated on day 5, and the plants allowed to grow for an additional 8 d such that leaves were theoretically producing PHB for a total of 13 d. Using this method quantifiable levels of PHB were detected both in the – RGG and + RGG conditions, whilst negligible PHB was detected in WT tissue (Figure 4D). This suggests that all three enzymes were active when targeted to the condensates.

Strikingly, localising a single or multiple enzymes to the biomolecular condensates resulted in 2.3- (citramalate) and 4.9- (PHB) fold higher product accumulation levels than achieved for purely cytosolic enzymes (P < 0.001 in both cases) (Figures 4C and D). Several reasons could explain this effect. Firstly, and as already explained, it is now known that localising enzymes to biomolecular condensates affects their kinetics (at least *in vitro*) both through mass action (i.e. through increased concentrations of enzymes and/or substrates) and spatial effects such as favourable orientation of the enzyme and its substrate [18]. Secondly, bringing multiple copies of enzymes forming part of a pathway together can encourage flux through that pathway by a phenomenon termed probabilistic metabolite channelling [62]; this would likely only affect flux through the multi-enzyme PHB pathway, however.

It is also possible that citramalate and/or PHB are accumulating inside the condensate, thus shielding them from any degradation and increasing final yield. Alternatively, production of PHB is known to be detrimental in a range of hosts possibly due to the high hydrophobicity of the PHB granules themselves [63]. By sequestering PHB in biomolecular condensates, this effect could be reduced, potentially increasing product accumulation by alleviating the strain on the host. To examine this, we looked at the localisation of PHB to ascertain whether it was retained within the condensates. PHB accumulates as insoluble granules which can be stained using Nile red, a widely used neutral lipid dye [64, 65]. Nile red-stained leaves 13 dpi with constructs targeting PHB enzymes to the condensates showed clear co-localisation of mClover3 (condensate scaffold) and Nile red signal (Figures 5A and B). This co-localisation was only seen in leaves expected to produce PHB: it was not seen 3 dpi when no PHB was detected (Figure S4) or when leaves were infiltrated with only the scaffold construct (Figures 5A and C). Given that Nile red also stains other neutral lipids, it is possible that PHB production somehow causes lipids to be sequestered inside biomolecular condensates and that these are then stained by Nile red, although this seems unlikely. Note that the apparent sequestration of PHB inside the condensates does not necessarily imply that citramalate is also enriched in the condensates, as condensate enrichment is known to be dependent on the physicochemical characteristics of the sequestered molecule [13].

**Figure 5:**
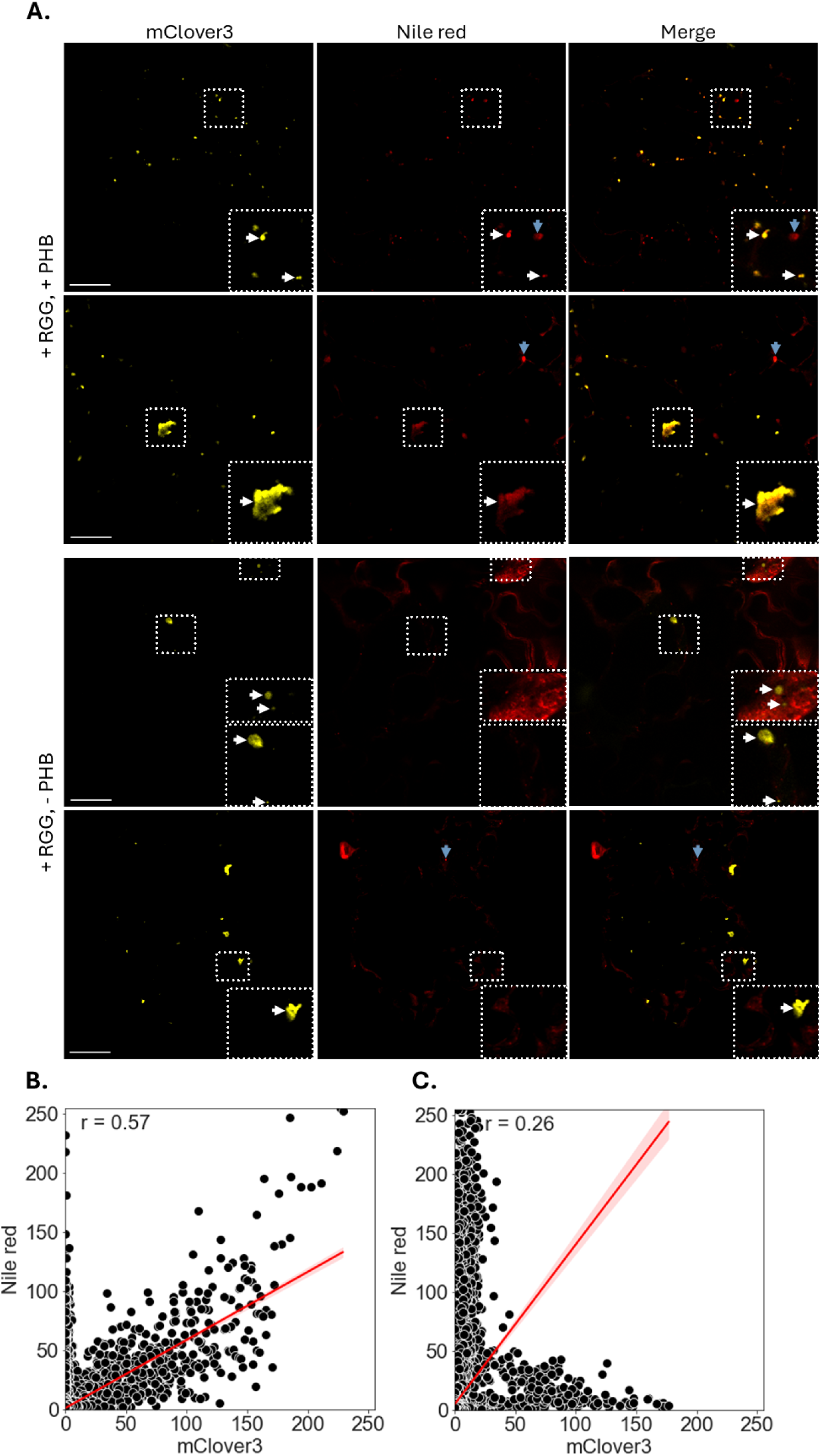
PHB localises to biomolecular condensates in N. benthamiana. (A) Co- localisation of mClover3 and Nile red signal in representative images of N. benthamiana leaf transiently expressing the RGG-containing scaffold (+ RGG) and the PHB-enzyme client (+ PHB) constructs (top), or the RGG-containing scaffold only (bottom). White arrows indicate a condensate, and blue arrows indicate a probable lipid droplet. Scale bar is 50 μm. (B) The relationship between Nile red and mClover3 signal in a leaf infiltrated with + RGG and + PHB. (C). The relationship between Nile red and mClover3 signal in a leaf infiltrated with + RGG only. In both (B) and (C) shaded area represents SEM, and the Pearsons correlation coefficient (r) is shown in the top-left corner.

It is unclear whether the PHB-producing enzymes are active inside the condensates, as opposed to just on the surface. Indeed, enzymatic activity and product production may be heterogeneously distributed throughout a condensate. Experiments involving the immobilisation of NanoLuc – a reporter enzyme that produces a luminescence signal upon reacting with its substrate, fumirazine – inside an RGG-based condensate *in vitro* demonstrated that the reaction was initially predominantly occurring at the water-condensate interface, and gradually reached the inside of the biomolecular condensate as the substrate diffused into the condensate [23].

If *Md*CMS and the PHB pathway enzymes are functional inside the condensates, it would follow that acetyl-CoA, H_2_O, pyruvate, and NADPH are able to diffuse into the condensates. No framework yet exists to predict which metabolites can access which condensates and why. It is likely that the physicochemical attributes of specific metabolites, including hydrophobicity and aqueous solubility, influence enrichment in a given condensate [13] although only a small number of condensates formed *in vitro* have thus far been tested. Being a substrate for both *Md*CMS and β-ketothiolase, it is possible that acetyl-CoA is being enriched in the condensates and that this is also contributing to the higher product yield. This would be significant given the yield-limiting nature of acetyl-CoA in many metabolic engineering applications [66].

### 2.5. The increased accumulation of condensate-targeted enzymes

As well as mass action kinetics, conformational effects, and selective sequestration of substrates and/or products, one final way in which condensate-targeting might be affecting product yield is through protein stabilisation. To determine whether this might be playing a role here we assessed the level of citramalate synthase (at 5 dpi) and of β-ketothiolase, acetoacetyl-CoA reductase, and PHB synthase (at 13 dpi). Confocal microscopy was used to determine the levels of mClover3 (as a proxy for the scaffold) and mCherry (as a proxy for the client[s]) in the – RGG and + RGG scenarios. There was a clearly higher level of both fluorescent proteins (up to 7.4-fold (PHB synthase), P < 0.01) in the + RGG scenarios and this was the same for mCherry fused to citramalate synthase and the enzymes of the PHB biosynthetic pathway (Figures 6A and S5).

**Figure 6:**
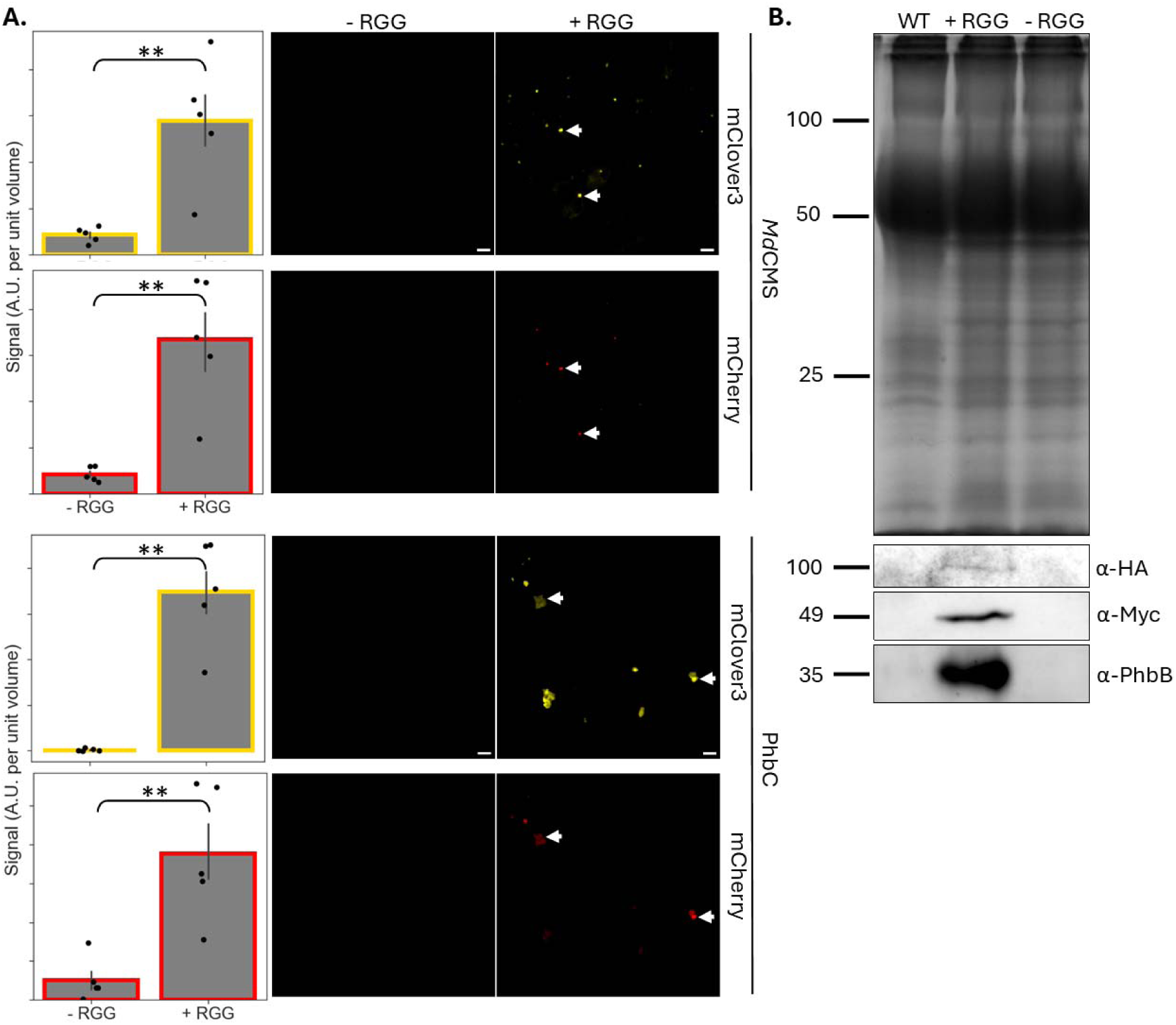
The accumulation of condensate-targeted enzymes. (A) Images were taken with identical acquisition settings and total mClover3 and mCherry signal used to provide a semi-quantitative measure of scaffold and client levels in each image. Biomolecular condensates are indicated using a white arrow. Results for PHB synthase (PhbC) are shown; for β-ketothiolase and acetoacetyl-CoA reductase see Figure S5. Error bars represent SEM; ‘***’ represents a P-value < 0.001, ‘**’ a P-value < 0.01, ‘*’ a P-value < 0.05, or ‘n.s.’ a P-value > 0.05 (Mann-Whitney U). n = 5. (B) SDS-PAGE (top) and western blot (bottom) of crude cell lysate from un-infiltrated leaves (WT) or leaves infiltrated with the RGG-containing (+ RGG) or non-RGG-containing (- RGG) version of the scaffold. Loading was normalised by starting mass of tissue. Bands on SDS-PAGE representing 100, 50, and 25 kDa are shown. An α-HA antibody was used to detect HA-tagged PHB synthase, an α- Myc antibody to detect Myc-tagged β-ketothiolase, and an α-PhbB antibody used to detect acetoacetyl-CoA reductase, with approximate expected sizes in kDa shown.

To make a more direct assessment of client enzyme accumulation, we used immunoblotting using an antibody against acetoacetyl-CoA reductase. Additional immunoblots were done against small epitope tags (Myc or HA) linked to the enzymes β-ketothiolase and PHB synthase (Figure 6B). β-ketothiolase, acetoacetyl-CoA reductase, and PHB synthase were detected in the crude lysate from leaves 13 dpi with the + RGG but no signal was detected in the – RGG condition for any of the proteins.

Hence the targeting of enzymes to biomolecular condensates results in an increase in enzyme accumulation. This result suggests that enzyme stabilisation could be partly responsible for the higher product yields achieved when enzymes are targeted to cytosolic biomolecular condensates. It is likely that the stabilisation effect is due to the protection afforded by the biomolecular condensates from proteolysis. In that context, it has been shown that recruitment of a synthetic degron-tagged mCitrine to a condensate reduces its degradation in *Escherichia coli* [67]. The striking increase in enzyme accumulation in *N. benthamiana* when targeted to biomolecular condensates has broad implications for protein overproduction and metabolic engineering. The effect is perhaps unsurprising given that proteins and RNA are known to be stabilised by recruitment to endogenous biomolecular condensates *in vivo* and this is used by the cell to control metabolism and development [12, 68].

### 2.6. Significance of findings for the development of the N. benthamiana biofactory

In this work we have shown that synthetic biomolecular condensates can be induced in transiently transformed *N. benthamiana* cells and that these can be functionalised by the targeting of a fluorescent protein, a single enzyme, or a three-step enzymatic pathway. Our results suggest that enzymes remain functional when targeted to the condensates, and that these condensates can sequester at least some types of products. The targeting of enzymes to biomolecular condensates results in a substantial increase in metabolic pathway product yield when compared to untargeted enzymes. We suggest that this is at least partly attributable to the significant increased accumulation of enzymes within transformed cells when targeted to biomolecular condensates *in planta*.

Controlled compartmentation of engineered pathways and protection from proteolysis are significant obstacles to the development of the *N. benthamiana* biofactory. Compartmentation has long been explored in metabolic engineering as a way to sequester a product from the rest of the cell either because it is toxic or because it is readily metabolised by the host. For instance, attempts at the heterologous production of cannabinoids [16] and squalene [69] in *N. benthamiana* have suffered from autotoxicity of products or intermediates and is an obstacle to high production levels. In addition, endogenous enzymes are known to affect the accumulation of a range of compounds in *N. benthamiana* including artemisinic acid and a range of terpenoids [70, 71]. The work presented here suggests that localising enzymes to biomolecular condensates could help address this challenge by facilitating product sequestration.

As well as product sequestration, this work has showed that another advantage of targeting enzymes to biomolecular condensates is increased product yield. Targeting of both a single enzyme and a three-enzyme heterologous pathway resulted in a 2.3- to 4.9-fold increase in product yield. Due to the substantive capital investments required to set up and run large-scale facilities for *N. benthamiana* agroinfiltration, it remains uncertain whether such platforms will be economically viable for the production of products that are not of exceptionally high commercial value [71]. Any increase in product yield would help mitigate this.

Plant-based protein production is arguably the best developed application of *N. benthamiana* agroinfiltration, and several protein products are currently on the market. However, recombinant protein production also suffers from low yield [72]. Here we showed that as well as an increase in product yield, targeting enzymes to biomolecular condensates increased their accumulation such that they were present at up to 7.4-fold higher levels. This was the case for four different enzymes suggesting it to be a universal feature of the system. This effect was maintained until at least 5 dpi for citramalate synthase, and 13 dpi for the three enzymes involved in PHB biosynthesis albeit with re-infiltration at 5 dpi. Note that the sequestration of proteins in biomolecular condensates may not be suitable for all targets; for instance, it may interfere with modifications such as N-glycosylation in the ER which are required for some pharmaceutical proteins [42].

## 3. Experimental procedures

### 3.1. Plant materials and growth

*N. benthamiana* was grown in a greenhouse with a 16:8 h light:dark cycle at 25 °C. Natural light was supplemented with ATTIS-7 LED grow lights (Plessey, London, UK) to achieve a light intensity of up to 200 μmol photons m^-2^ s^-1^. Plants were grown in 11-cm square pots on soil composed of a 3:1 ratio of modular seed growing compost and pro fine vermiculite (both Sinclair Pro, Cheshire, UK) and were regularly watered. 0.4 g L^-1^ Exeptor (ICL, Ipswich, UK) was used as an insecticide.

### 3.2. Molecular cloning and genetic construct fabrication

Gene parts used in genetic construct assembly are listed in Table S1. Most gene parts were synthesised by Twist Bioscience (San Francisco, California, USA) and codon optimised for *N. benthamiana* using the in-house submission tool. The Gateway vector pK7WG2 [47] was used for expression. Plasmids were designed and assembled *in silico* using SnapGene (version 7.1.0, GSL Biotech, San Diego, California, USA).

DNA amplification using polymerase chain reaction (PCR) was carried out using Phusion High-Fidelity DNA Polymerase (Thermo Fisher Scientific, Waltham, Massachusetts, USA) according to the manufacturer’s instructions. Primer information can be found in Table S2. PCR fragments were assembled for transformation using the NEBuilder HiFi DNA Assembly Cloning Kit (New England Biolabs, Hitchin, UK). Constructs were assembled into the pTWIST-Entr Gateway cloning vector, and the Gateway LR Clonase II Enzyme Mix (Thermo Fisher Scientific, Waltham, Massachusetts, USA) used to move the constructs into pK7WG2 according to the manufacturer’s instructions.

Cloning was undertaken using α-select Silver Chemically Competent *E. coli* cells (Meridian Bioscience, Cincinnati, Ohio, USA) using a standard heat shock transformation protocol. At least 5 colonies per construct were screened using diagnostic restriction digestion with an appropriate FastDigest restriction enzyme (Thermo Fisher Scientific, Waltham, Massachusetts, USA) followed by Sanger sequencing (Source Bioscience, Nottingham, UK) and full plasmid sequencing (Plasmidsaurus, Eugene, Oregon, USA). Validated sequences were transformed into *Agrobacterium tumefaciens* strain GV3101 (pMP90) [73] using a standard freeze-thaw protocol. For all plasmids see Table S3.

### 3.3. Plant transformation

*N. benthamiana* plants between 4 and 5 weeks old were transiently transformed by injecting *A. tumefaciens* in agroinfiltration buffer (10 mM MES monohydrate, 10 mM MgCl_2_, 100 μM acetosyringone, pH 5.7) at OD_600_ of 0.5 or 1.0 into the underside of mature leaves using a 1 mL blunt-ended syringe. To increase expression levels, constructs were co-infiltrated with the RNA silencing suppressor P19-carrying strain (Emma Watts, University of Oxford, UK) [74] in a 1:1 ratio. During co-infiltration experiments, *A. tumefaciens* strains were mixed in equal ratios and at the same OD_600_ in agroinfiltration buffer. In some cases, to ensure adequate expression levels, plants were re-infiltrated 5 d after the first infiltration event with an identical *A. tumefaciens* solution.

### 3.4. Confocal laser scanning microscopy

Leaf epidermal samples of *N. benthamiana* were imaged using a Leica SP5 confocal microscope (Leica Microsystems, Wetzlar, Germany). Leaf disks were excised from live leaves and mounted with the abaxial side upwards in perfluorodecalin (Merck, Darmstadt, Germany). Images were captured with an HC PL APO 20× water immersion objective lens or an HC PL APO CS2 63× water immersion objective lens (both Leica Microsystems, Wetzlar, Germany) at 512×512 px resolution with 4-line averaging and gain adjusted to maximize signal whilst avoiding saturation. Saturation was evaluated using the LAS AF Application Software (Leica Microsystems, Wetzlar, Germany) prior to image acquisition. An excitation wavelength of 488 nm for mClover3 or 594 nm for mCherry was used, with emission collected at 510-540 nm or 600-630 nm, respectively. When collecting z-stacks 1-line averaging was used to reduce the delay between images.

For the nuclear staining, FRAP, concentration-dependent condensate formation, and protein stability experiments, samples were imaged using a Zeiss LSM 880 confocal microscope with a LD LCI Plan-Apochromat 25x/0.8 mm or a C-Apochromat 40x/1.2 mm water immersion objective lens (both Carl Zeiss AG, Oberkochen, Germany).

To stain nuclei the underside of agroinfiltrated *N. benthamiana* leaves were injected with 100 ng mL^-1^ 4’,6-diamidino-2-phenylindole (DAPI) in phosphate buffered saline (137 mM NaCl, 2.7 mM KCl, 10 mM Na_2_HPO_4_, 1.8 mM KH_2_PO_4_, pH 7.4) using a 1 mL blunt-ended syringe. Leaf samples were excised and mounted as previously described and images captured after 5 min with excitation at 405 nm (DAPI), 488 nm (mClover3), and 561 nm (mCherry), and emission collected at 430-460 nm (DAPI), 510-540 nm (mClover3), and 600-630 nm (mCherry).

For FRAP experiments, circular regions encompassing an entire or part of a compartment were bleached using the 488 nm laser at 100 % intensity for 100 iterations. The minimum size of bleached area was limited by the size of the laser. Images were taken for five pre- and 70 post-bleach scans with an excitation of 488 nm and emission collected at 510-540 nm with the delay between timeseries images minimized. Images were kept for analysis if signal was reduced by at least 50 % in the bleached area, and laser intensity during imaging was decreased to avoid any photobleaching as revealed by monitoring signal in a non-bleached area.

For the concentration-dependence and protein stability experiments, mClover3 only or mClover3 and mCherry signal, respectively, were captured as z-stacks with the emission and excitation settings already described. All images within the same experiment were acquired using identical settings including pixel size, objective, light path, laser power, gain offset, frame size, zoom level, scan speed and pinhole size. Extra care was taken to avoid any saturated pixels by using the ZEN Microscopy Software (Carl Zeiss AG, Oberkochen, Germany) prior to image acquisition.

For the PHB staining experiment, samples were imaged using a Leica Stellaris confocal microscope (Leica Microsystems, Wetzlar, Germany). Images were captured with an HC PL APO 20× water immersion objective lens or an HC PL APO CS2 63× water immersion objective lens (both Leica Microsystems, Wetzlar, Germany). To stain PHB granules the underside of agroinfiltrated *N. benthamiana* leaves were injected with 2 μg mL^-1^ Nile red in 50 mM piperazine-1,4-bis(2-ethanesulfonic acid) (pH 7) using a 1 mL blunt-ended syringe. Leaves were stained for 30 min and were excised and mounted as previously described. Images were captured sequentially with excitation at 480 nm (mClover3) and 540 nm (Nile red), and emission collected at 510-540 nm (mClover3) and 550-630 nm (Nile red).

### 3.5. Confocal image analysis

To calculate condensate size and number, a z-stack image spanning a single cell was captured on a Leica SP5 confocal microscope (Leica Microsystems, Wetzlar, Germany) using the settings for mClover3 already presented and was processed using ImageJ (version 1.54, U.S. National Institutes of Health, Bethesda, USA) [75] and the 3D Objects Counter plugin package (version 1.5.1;[76]). The threshold for object detection and size was manually adjusted for each image based on visual inspection to detect all condensates.

FRAP data was analysed using ImageJ and the Stowers ImageJ Plugins (version 2; Jay Unruh, Stowers Institute for Medical Research, Kansas City, Missouri, USA). Briefly, a circular bleached area was selected on a time course image and average pixel intensity quantified over time. For comparison across images pixel intensity was normalised by setting the maximum of each recovery curve to 1 and the minimum to 0.

For the concentration-dependence experiments, 3D (*x,y,z*) *.czi images were loaded into MatLab® (Mathworks, Natick, Massachusetts, United States of America) using the Bioformats reader [77], median filtered using a 5×5×3 pixel kernel and the background subtracted using the mode for the full 3D image. An intensity-weighted histogram was calculated and fit with a 2,3, or 4 component Gaussian mixture model to identify the background, nucleoplasm and dense phase(s) if present. Upper and lower bounds were constrained to ensure robust fits and the model with the highest adjusted rsquare selected. The nucleus was segmented from background using the mean+3×SD from the first Gaussian component, whilst the dense phase was segmented from the nucleoplasm using the mean+3×SD of the second component. Compartment volumes were calculated from the number of segmented pixels, and relative concentrations from the average pixel intensity in each compartment.

For the semi-quantitative analysis of protein stability, data analysis was undertaken in ImageJ. Sum projections were generated for z-stacks of individual channels, and the background subtracted by calculating the mean intensity in a 10×10 pixel region of interest. The ImageJ Analyze feature was used to calculate an integrated intensity for each image which was normalised by the number of stacks.

### 3.6. Protein purification and analysis

For soluble protein purification leaf fragments were flash frozen and homogenised in a TissueLyser Reaction Tube Holder cooled to -80°C using a TissueLyser II (Qiagen, Hilden, Germany) for 5 min at 25 Hz and resuspended in a 10-fold excess (v/w) of cold grinding buffer (50 mM NaH_2_PO_4_, 150 mM NaCl, 50 mM ascorbic acid, 0.6 % [w/v] PVPP-40, 0.4 % [w/v] bovine serum albumin, 5 % [v/v] glycerol, 1 % [v/v] TWEEN-20, 1 mini tablet of Pierce Protease Inhibitor [Thermo Fisher Scientific, Waltham, Massachusetts, USA], pH 8.0). The lysate was sonicated for 10 min using a Grant Ultrasonic Bath XB3 (Keison Products, Chelmsford, UK) and centrifuged for 15 min at 3000×g at 4°C and the supernatant collected.

For Western blots, proteins were fractionated using sodium-dodecyl-sulphate polyacrylamide gel electrophoresis (SDS-PAGE) and transferred to a nitrocellulose membrane using standard methods. To detect mClover3 and mCherry, α-GFP and α-mCherry antibodies (both Proteintech Europe, Manchester, UK) were used, respectively, at a concentration of 1 μg mL^-1^. Proteins containing c-Myc and HA tags were detected with commercial antibodies (Abcam, Cambridge, United Kingdom and Merck, Darmstadt, Germany, respectively [at 1 μg mL^-1^]). Acetoacetyl-CoA reductase was detected using an α-PhbB antibody (1:500 dilution) [78]. Bound antibodies were detected using a goat α-rabbit IgG antibody linked to horseradish peroxidase (HRP) (Merck, Darmstadt, Germany) and the EZ-ECL Chemiluminescence Detection Kit (Biological Industries, Kibbutz Beit-Haemek, Israel).

### 3.7. Polymer and metabolite analysis

The extraction of PHB was based on a method outlined in [79]. 20-250 mg of plant tissue was collected and flash frozen with a metal bead in LiN_2_. The tissue was homogenised in a TissueLyser Reaction Tube Holder cooled to -80 °C using a TissueLyser II (Qiagen, Hilden, Germany) for 5 min at 25 Hz, transferred to a glass tube and extracted in 4 mL methanol for 1 h at 55 °C and subsequently extracted twice in 4 mL ethanol for 1 h at 55 °C. This was followed by the addition of 500 μL chloroform, 1.7 mL ethanol, 0.2 mL HCl, and 2 μL internal standard 3-hydroxy-valerate-methyl-ester to each sample, which was incubated for 4 h at 100 °C, resulting in the formation of 3-hydroxybutyrate ethyl ester. Lastly 4 mL of 0.9 M NaCl was added, the tubes vortexed, and the phases separated by centrifugation at 4,000×g for 10 min at 4 °C. The upper aqueous phase was removed and 25 μL of the lower phase was derivatised with 75 μL of N-methyl N-trimethylsilyltrifluoroacetamide for 30 min at room temperature with gentle shaking.

For citramalate extraction a method developed by Lisec et al was used [80]. 20-200 mg of plant tissue was flash frozen with a metal bead in LiN_2_ and homogenised in a TissueLyser Reaction Tube Holder cooled to -80 °C using a TissueLyser II (Qiagen, Hilden, Germany) for 2 min at 30 Hz. After homogenisation 400 μL pre-cooled methanol was added, the tube was vortexed, followed by addition of 60 μL 0.1 mg mL^-1^ norvaline as the internal standard. Samples were incubated at 70 °C for 15 min with shaking, before being centrifuged for 10 min at 20,000×g. The supernatant was transferred to another tube and 250 μL chloroform and 500 μL water added. Each tube was vortexed for 15 sec and then centrifuged for 15 min at 1700×g. Two samples of 100 μL were taken from each tube to generate duplicate samples to be dried down under vacuum. To derivatise the dried samples, 25 μL pyridine was added and the samples incubated at 37 °C with shaking at 950 RPM for 90 min followed by the addition of 35 μL *tert*-butyldimethylsilane and incubation at 65 °C with shaking at 950 RPM for 1 h.

0.25 μL (PHB quantification) or 0.5 μL (citramalate quantification), was injected in splitless mode at 230 °C into an Intuvo 900 gas chromatography mass spectrometer (Agilent Technologies, Santa Clara, USA) fitted with a 15 m × 250 μm × 0.25 μm DB-5ms Ultra Inert Intuvo GC capillary column module (Agilent Technologies, Santa Clara, USA). For PHB quantification, chromatography was achieved by holding the temperature at 60 °C for 5 min followed by heating at a rate of 7.5 °C min^-1^ to 200 °C resulting in a run time of 24 min. For the quantification of citramalate the temperature was held at 90 °C for 3 min followed by heating at a rate of 7.5 °C min^-1^ to 250 °C resulting in a run time of 24 min. For both PHB and citramalate quantification, an Agilent 5977B mass spectrometer (Agilent Technologies, Santa Clara, USA) was used in single ion monitoring (SIM) mode to detect the 189 and 203 *m/z* fragment ions for PHB and 3-hydroxy-valerate-methyl-ester respectively, and the 433 and 288 *m/z* fragment ions for citramalate and norvaline quantification respectively. Samples were injected in triplicate and average values taken forward for analysis. In all cases the MassHunter MS Quantitative Analysis software (Agilent Technologies, Santa Clara, USA) was used for peak identification and integration with the help of the National Institute of Standards and Technology (NIST, Gaithersburg, USA) mass spectrum library (version 2.0). PHB:3-hydroxy-valerate-methyl-ester and citramalate:norvaline standard curves were generated using pure PHB and citramalate standards (Merck, Darmstadt, Germany) and used for absolute quantification.

### 3.8. Data analysis and statistics

Statistics and data visualisation were performed using Python (version 3.8.18, Python Software Foundation, https://www.python.org) in a Jupyter Notebook [81]. A Mann Whitney U test [82] was used to determine differences between two groups and was implemented using the SciPy package [83]. Figures were prepared using the seaborn [84] and matplotlib [85] packages.

## Author contributions

A.L.L.B. and L.J.S. planned and designed the research. A.L.L.B. and A.W.B. carried out the experimental work. A.L.L.B., A.W.B., and M.D.F. analysed the data. A.L.L.B. wrote the first draft. All authors approved the final version of the manuscript.

## Acknowledgments

The authors thank the Interdisciplinary Bioscience Doctoral Training Programme at the University of Oxford for a full DPhil (PhD) studentship which was funded through the Biotechnology and Biological Sciences Research Council (funding number BB/M011224/1). The Gateway expression vector pK7WG2 and the mClover3 coding sequence were gifts from Francesco Licausi (University of Oxford, UK). The α-PhbB antibody was a gift from Yves Poirier (Université de Lausanne, Lausanne, Switzerland). We thank Pedro Bota for training and maintenance of the GC-MS.

---

**Figure S1:**
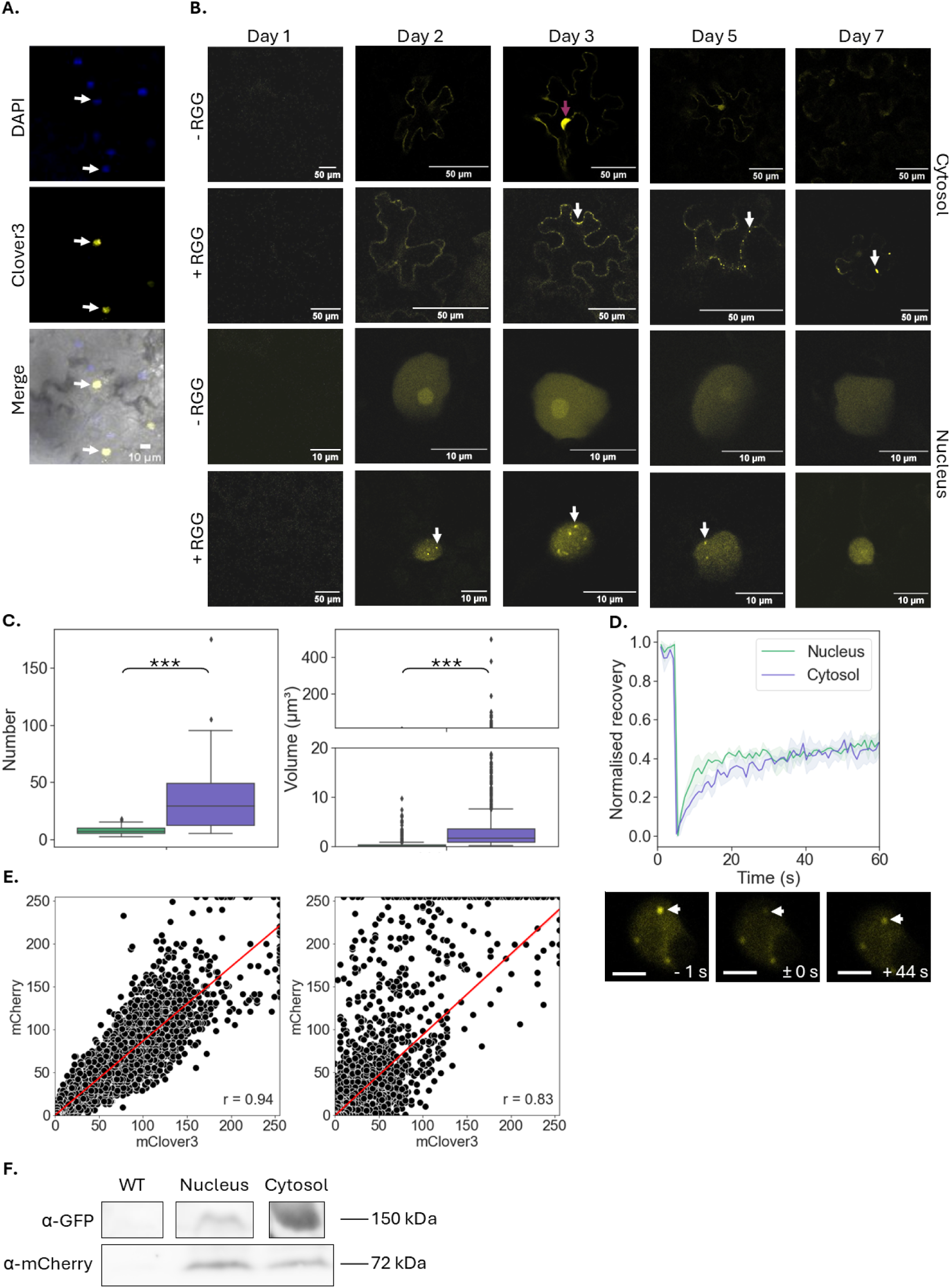
Further characterisation of synthetic biomolecular condensates in N. benthamiana. (A) DAPI-staining co-localises with mClover3 signal when an NLS is included, indicating that mClover3 gets targeted to the nucleus. (B) Timecourse of expression over 7 dpi in nuclei and cytosol of tissue infiltrated with the RGG-containing or no-RGG-containing version of the scaffold. mClover3 signal is shown. White arrows indicate condensates. Purple arrow indicates a nucleus; the presence of signal in the nuclei of cells overexpressing a fluorescent protein is common. (C) A comparison of the number per cell (left) and volume (right) of cytosolic (blue) and nuclear (green) biomolecular condensates. The distribution is plotted using a box and whisker plot, where the box shows the mean and the interquartile range. The whiskers extend to values above or below 1.5 × the interquartile range. Points either above or below this value are shown as diamonds. ‘***’ represents a P-value < 0.001, ‘**’ a P-value < 0.01, ‘*’ a P-value < 0.05, or ‘n.s.’ a P-value > 0.05 (Mann-Whitney U). n = 21-54 (number) or 400-880 (volume). (D) Normalised FRAP curves for nuclear and cytosolic biomolecular condensates (top), and example of biomolecular condensate bleaching in a nucleus (bottom). n = 4-5. Arrow indicates bleached biomolecular condensate. Pre (-) or post (+) bleaching time indicated. Scale bar is 10 μm and shaded area represents SEM. (E) The relationship between mCherry and mClover3 signal in a leaf expressing the nuclear (left) or cytosolic (right) version of the scaffold. Shaded area represents SEM, and the Pearsons correlation coefficient (r) is shown in the bottom-right corner. (F) Western blot for the detection of mClover3 (α-GFP) and mCherry in crude leaf extract of agroinfiltrated N. benthamiana leaves 5 dpi.

**Figure S2:**
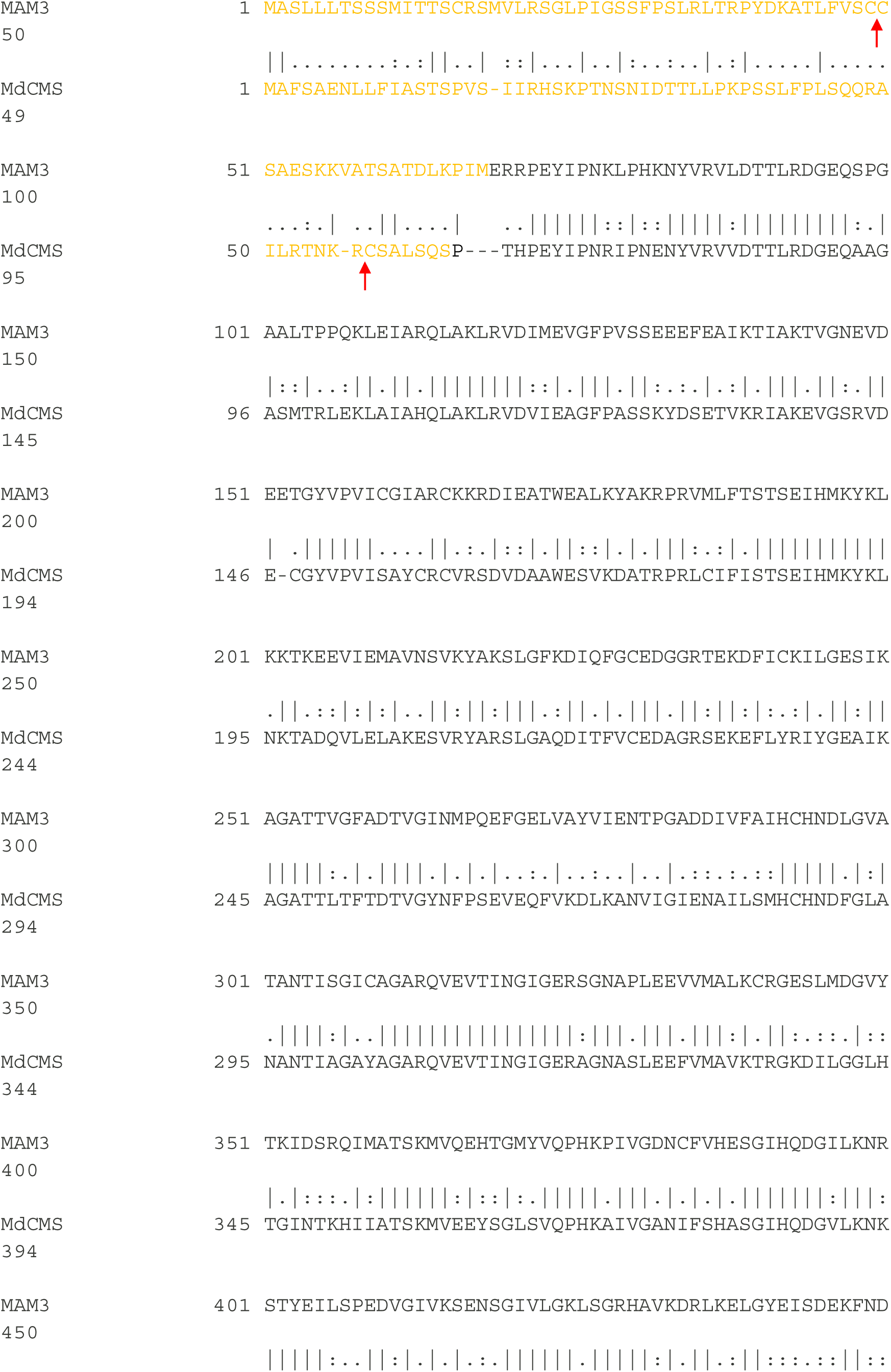
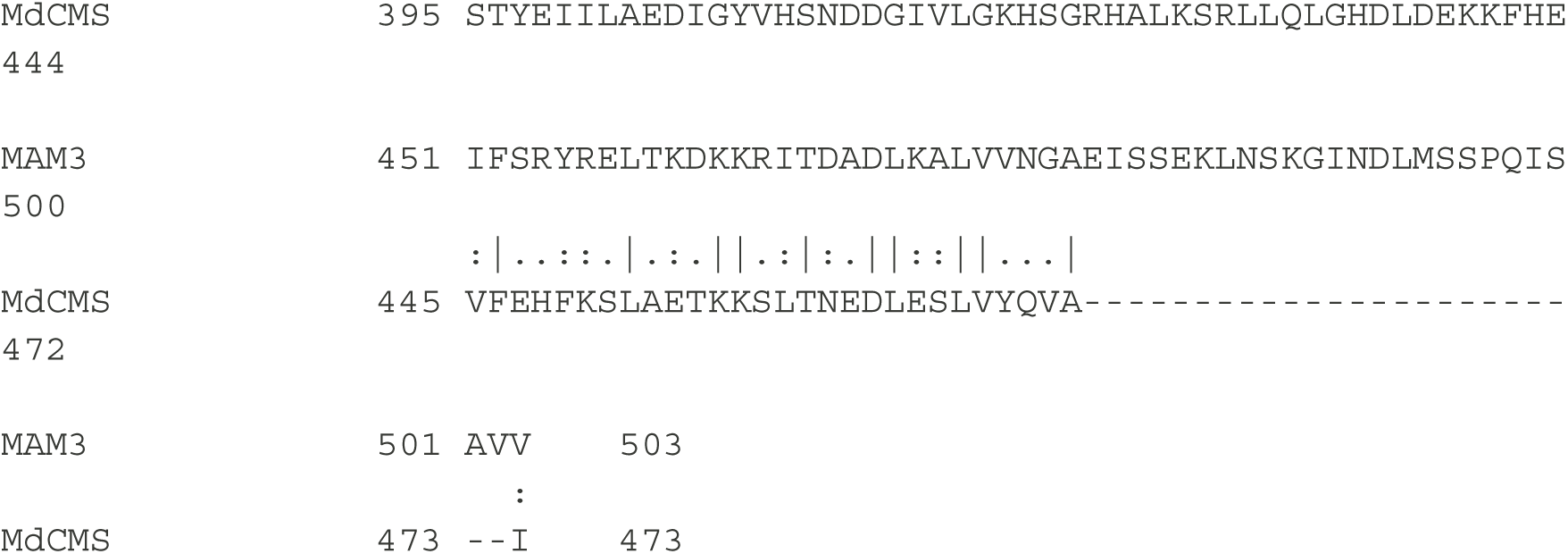
Alignment between MdCMS and closely related methylthioalkylmalate synthase 3 (MAM3) from A. thaliana used to determine whether MdCMS contains a chloroplast transit peptide. The alignment analysis was performed using EMBOSS Stretcher from EMBL-EBI [90]. MAM3 is known to be chloroplast localised and has a long unstructured region on its N-terminus which is predicted to encode a chloroplast transit peptide (see UniProt accession Q9FN52). In the predicted structure for MdCMS, the N-terminus is also unstructured when analysed using AlphaFold [91] and so we hypothesized that MdCMS also has an N-terminal chloroplast transit peptide. TargetP-2.0 [92] was used to predict the presence of a N-terminal chloroplast transit peptide with high probability, including a putative cleavage site, in the MdCMS protein sequence. N-terminal unstructured region indicated in orange and predicted cleavage sites shown using a red arrow.

**Figure S3:**
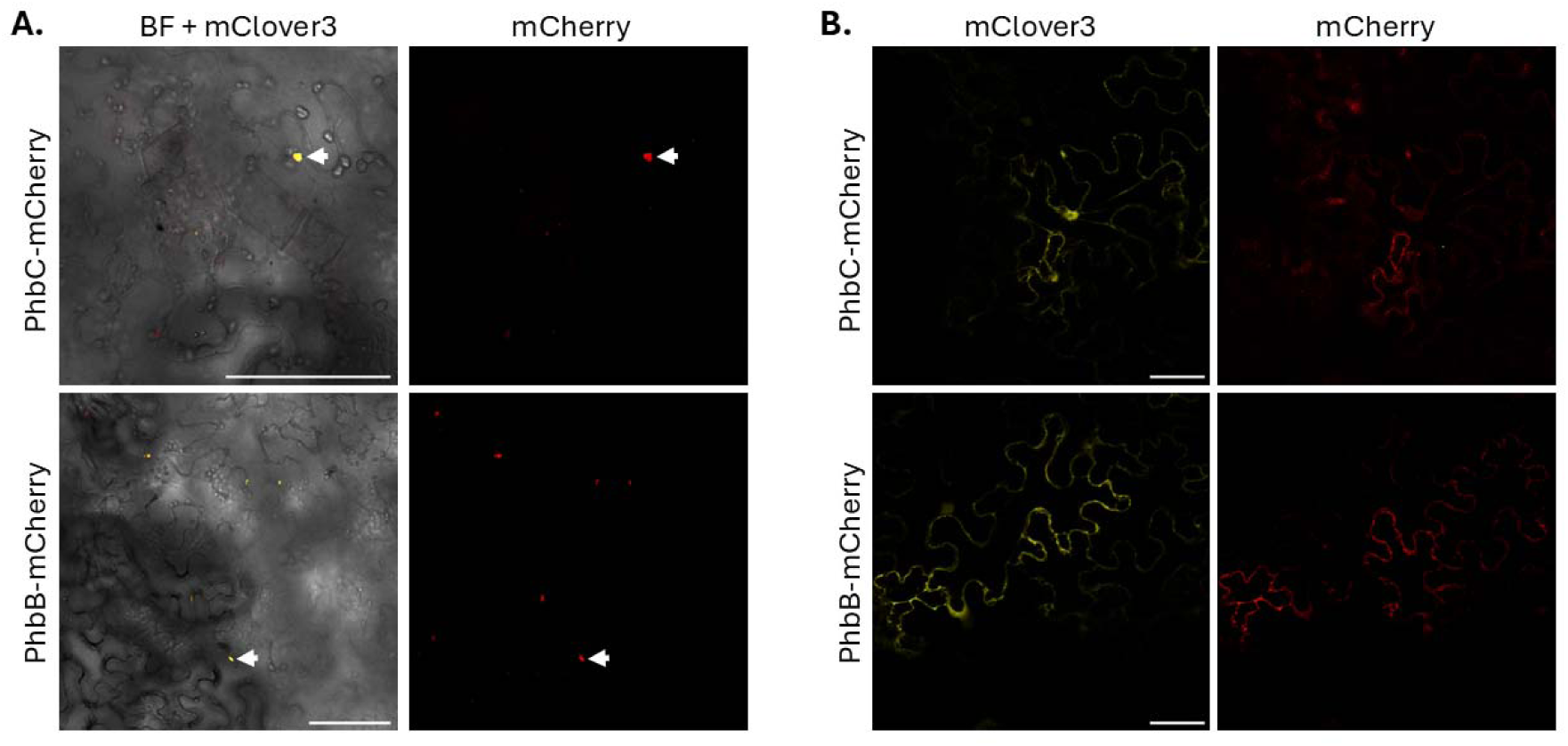
Further characterisation of PHB-functionalised condensates. (A) Co-localisation of mClover3 and mCherry-fused acetoacetyl-CoA reductase (PhbB), and PHB synthase (PhbC) in biomolecular condensates. White arrow indicates a condensate. (B) The removal of the RGG domain ablates biomolecular condensate formation and results in diffuse mClover3 and mCherry cytosolic signal. Scale bar in both panels is 100 μm.

**Figure S4:**
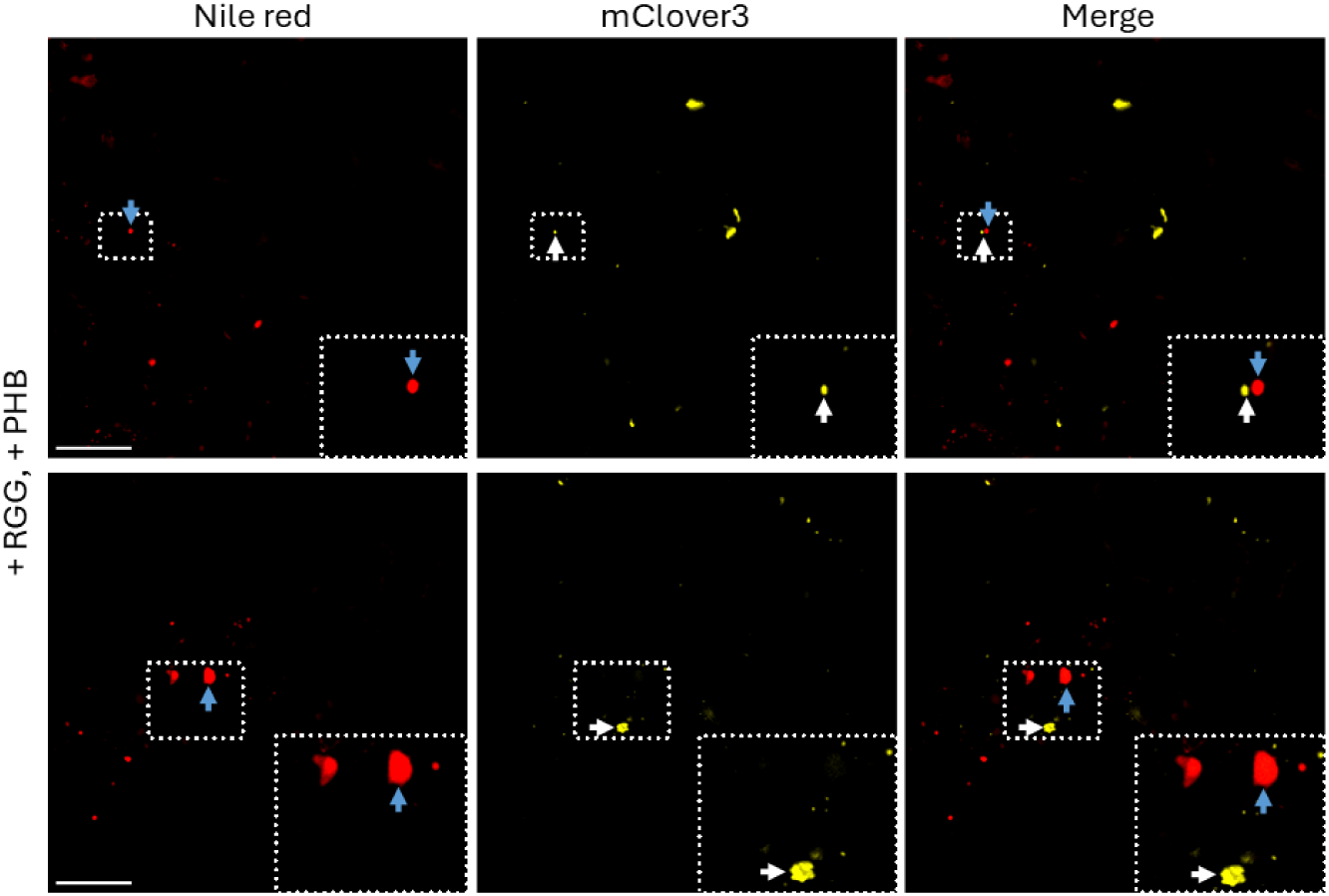
Absence of co-localisation of mClover3 and Nile red signal in representative images of N. benthamiana leaf transiently expressing the RGG-containing scaffold (+ RGG) and the PHB-enzyme client (+ PHB) constructs 3 dpi. White arrows indicate a condensate, and blue arrows indicate a probable lipid droplet. Scale bar is 50 μm.

**Figure S5:**
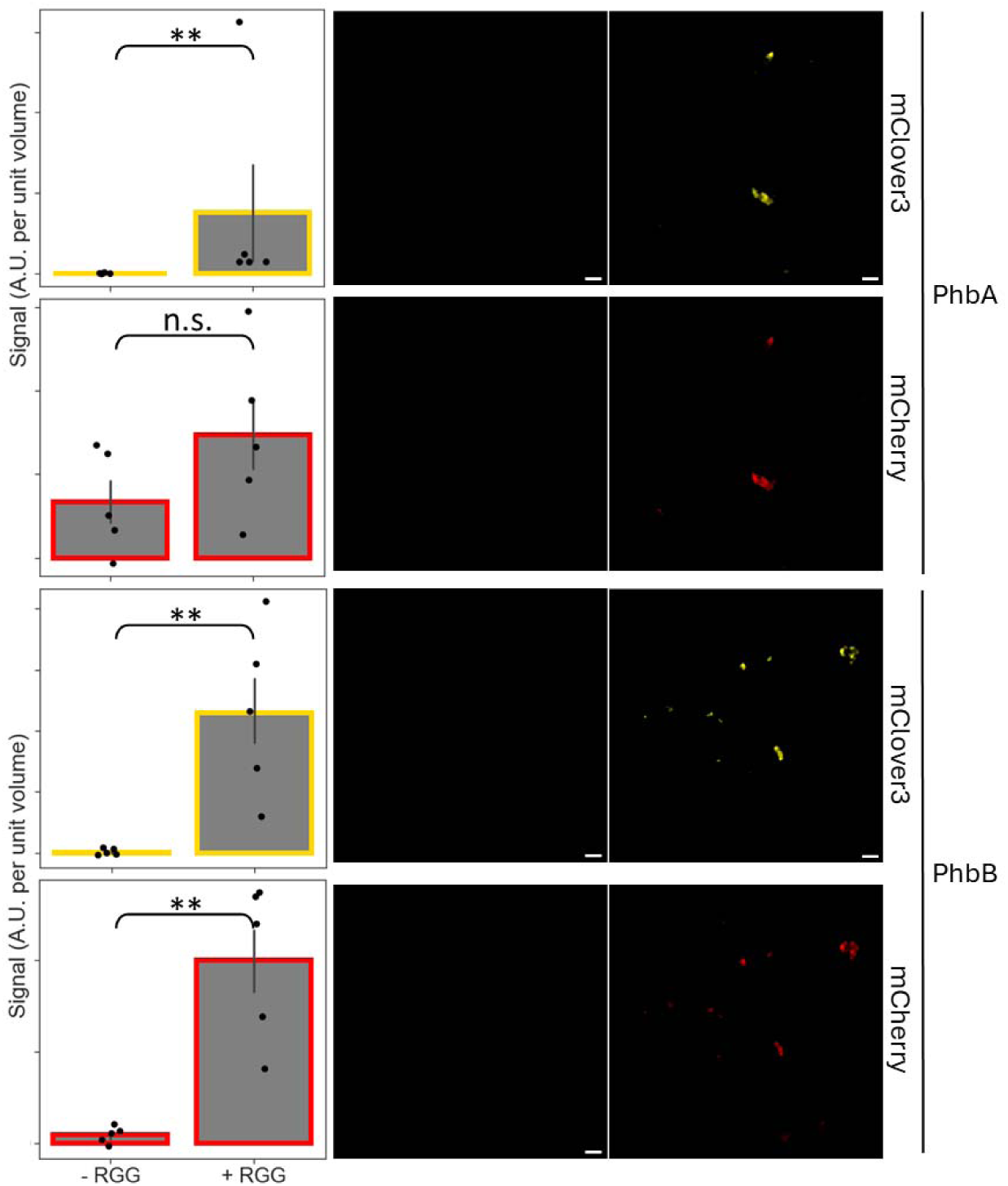
Semi-quantitative analysis of scaffold and client levels in each image, shown here for β-ketothiolase (PhbA) and acetoacetyl-CoA reductase (PhbB). Error bars represent SEM; ‘***’ represents a P-value < 0.001, ‘**’ a P-value < 0.01, ‘*’ a P-value < 0.05, or ‘n.s.’ a P-value > 0.05 (Mann-Whitney U). n = 5.

**Table S1:**
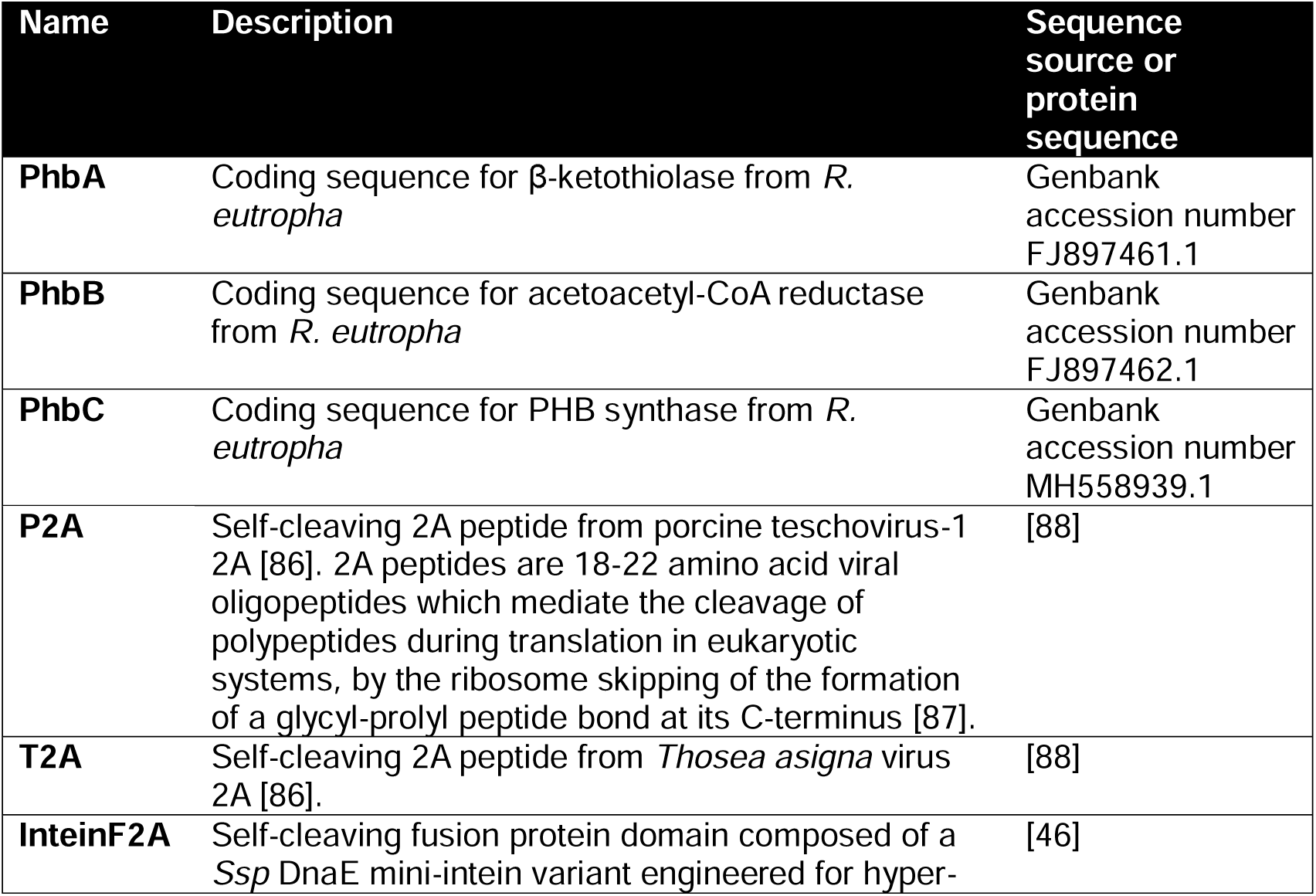
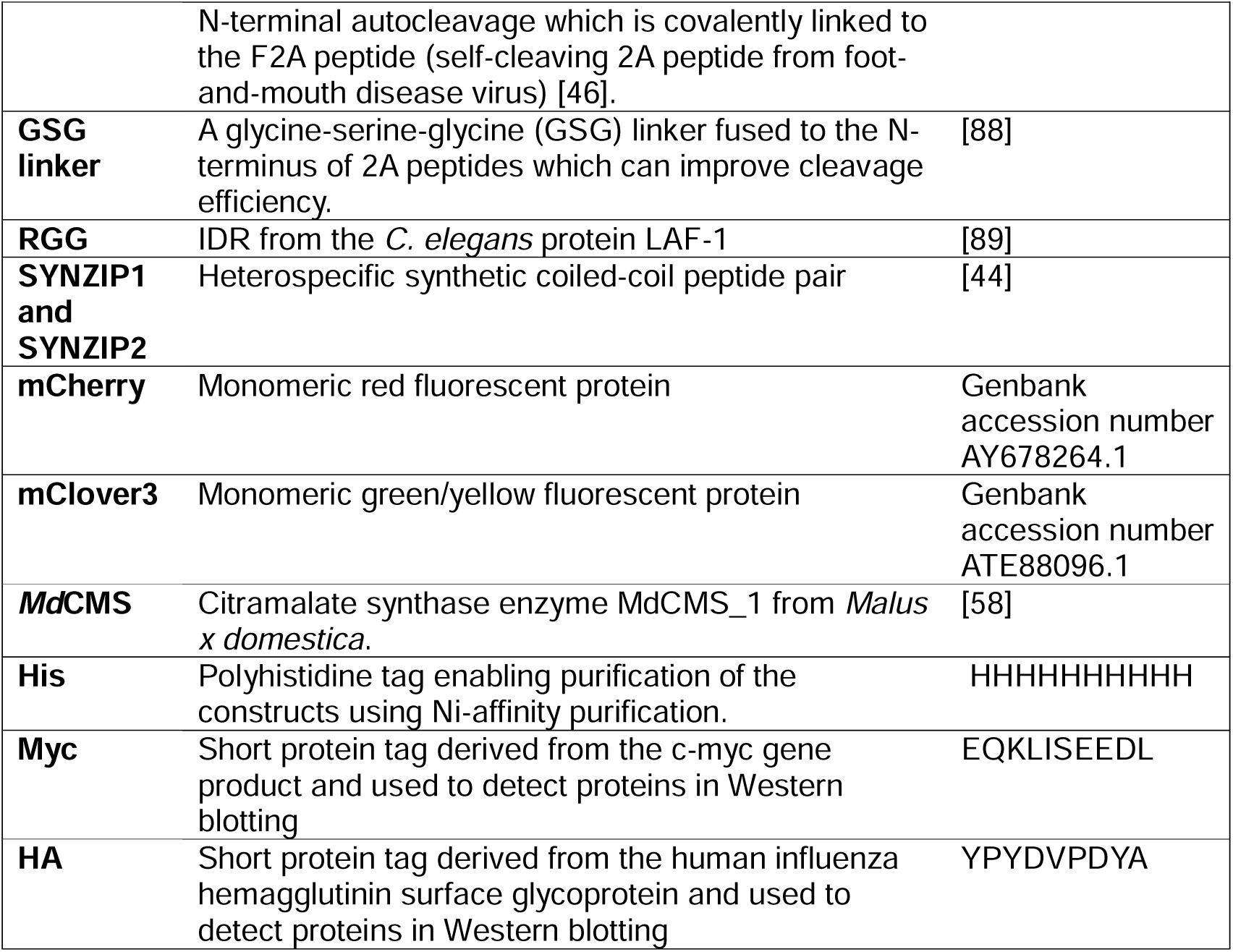
Gene parts used in this study.

**Table S2:**
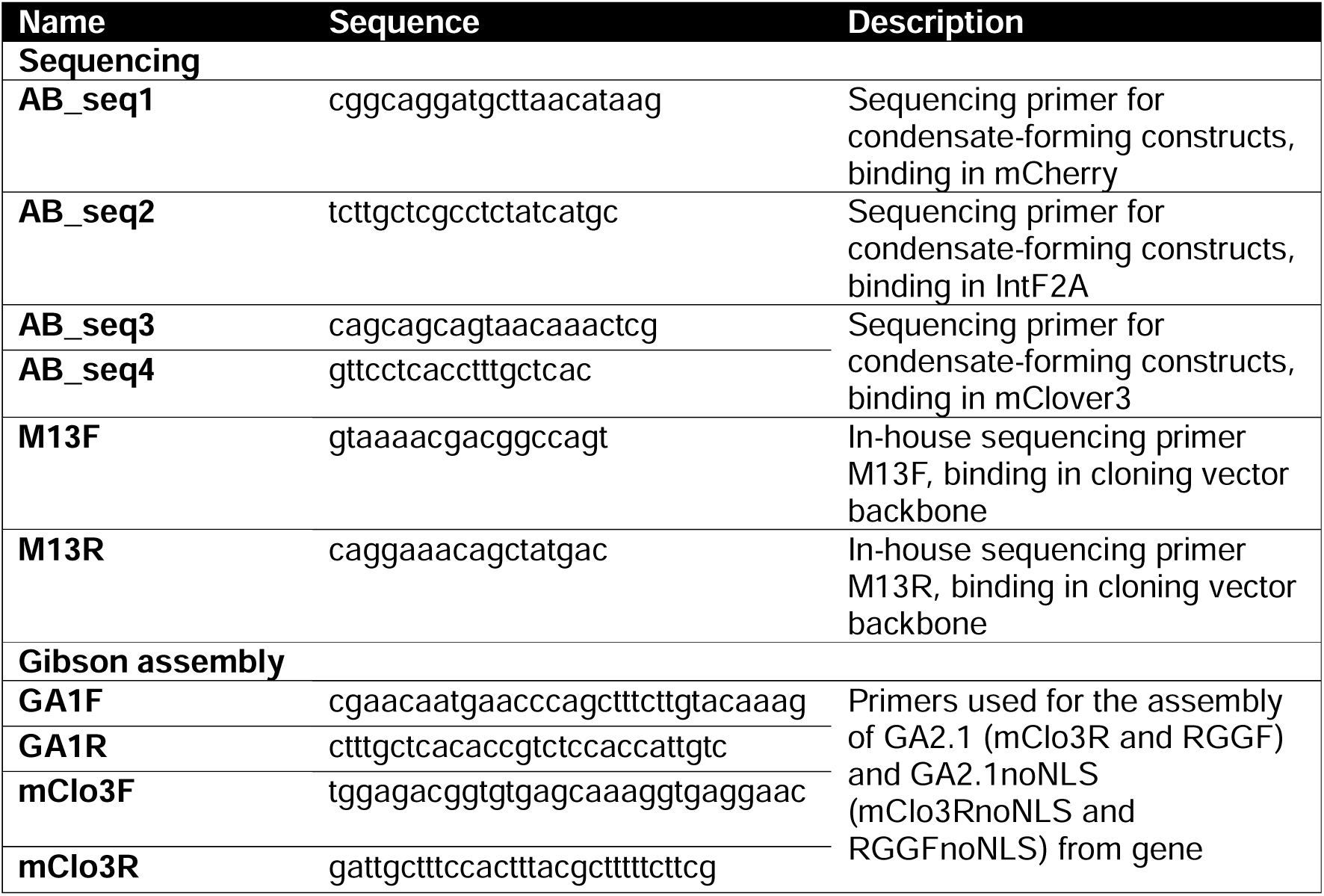
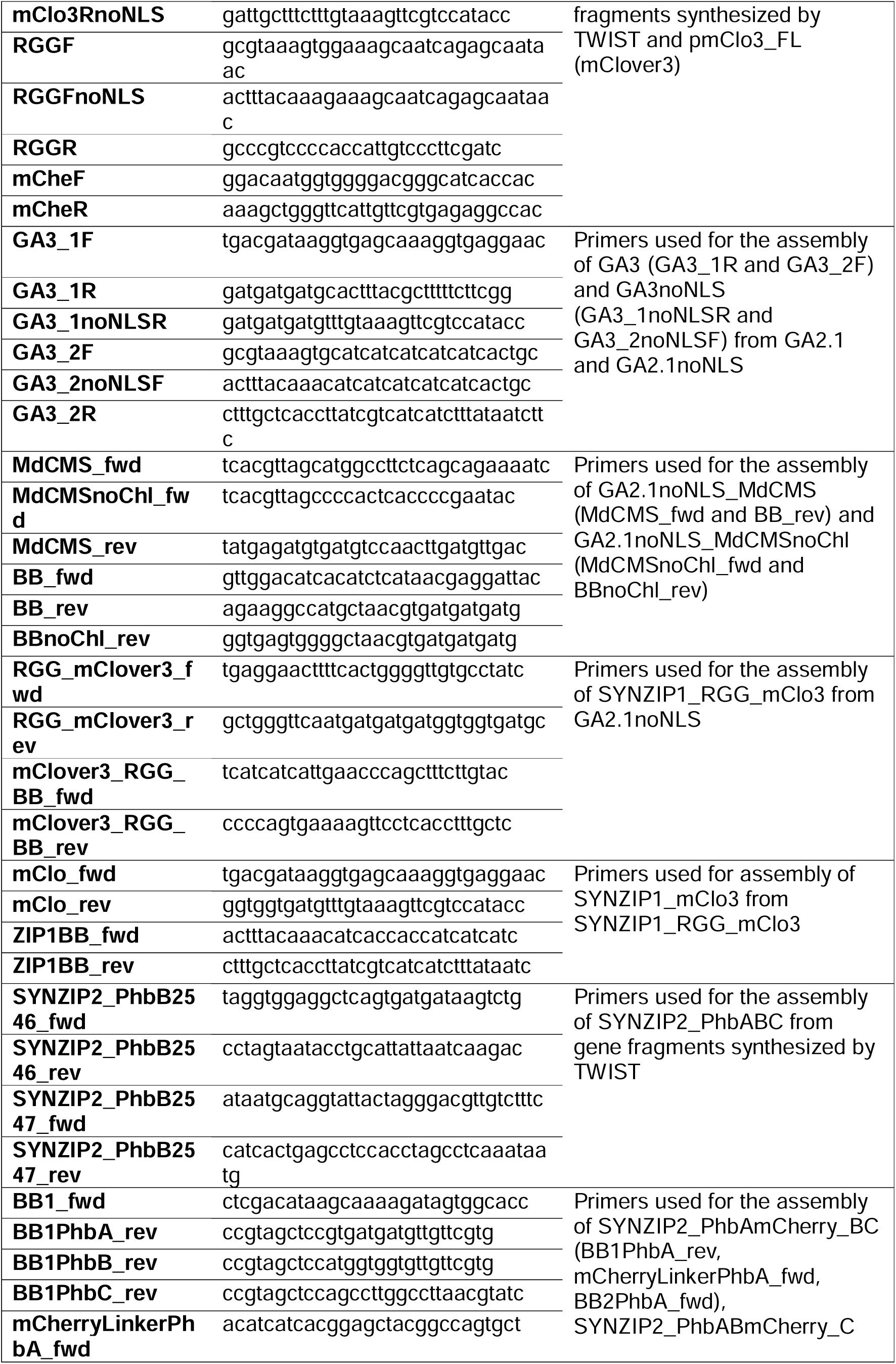
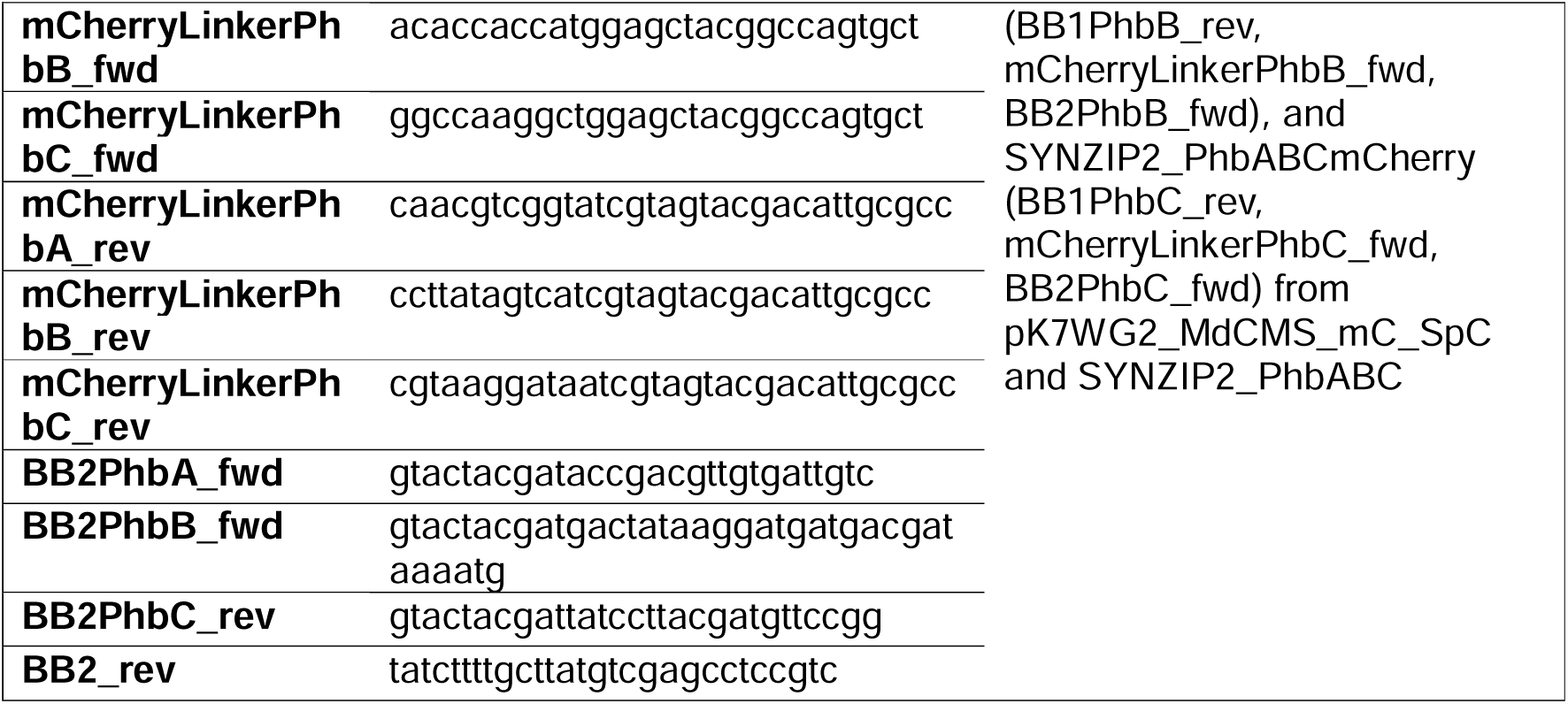
Primers used in this study.

**Table S3:**
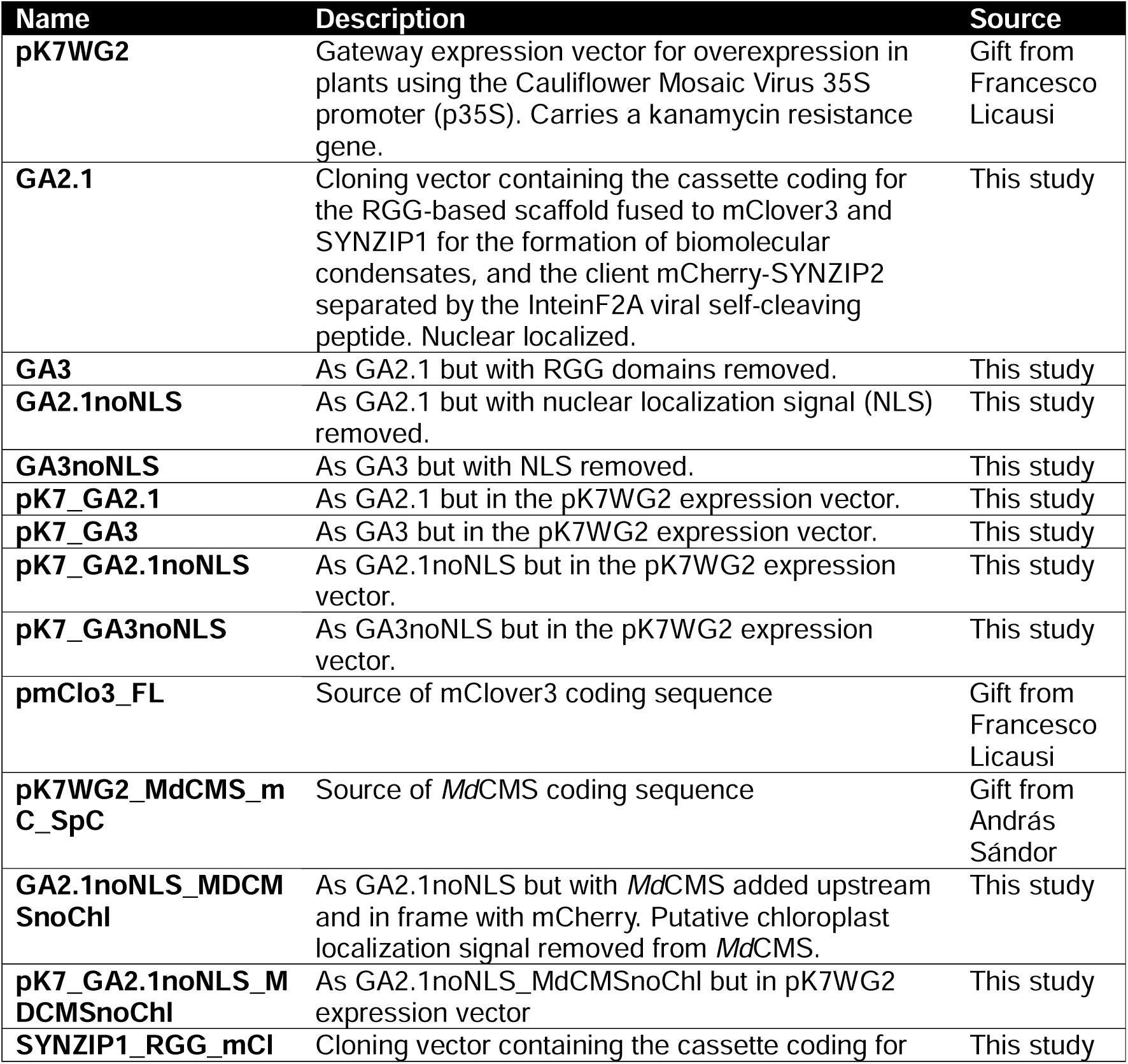
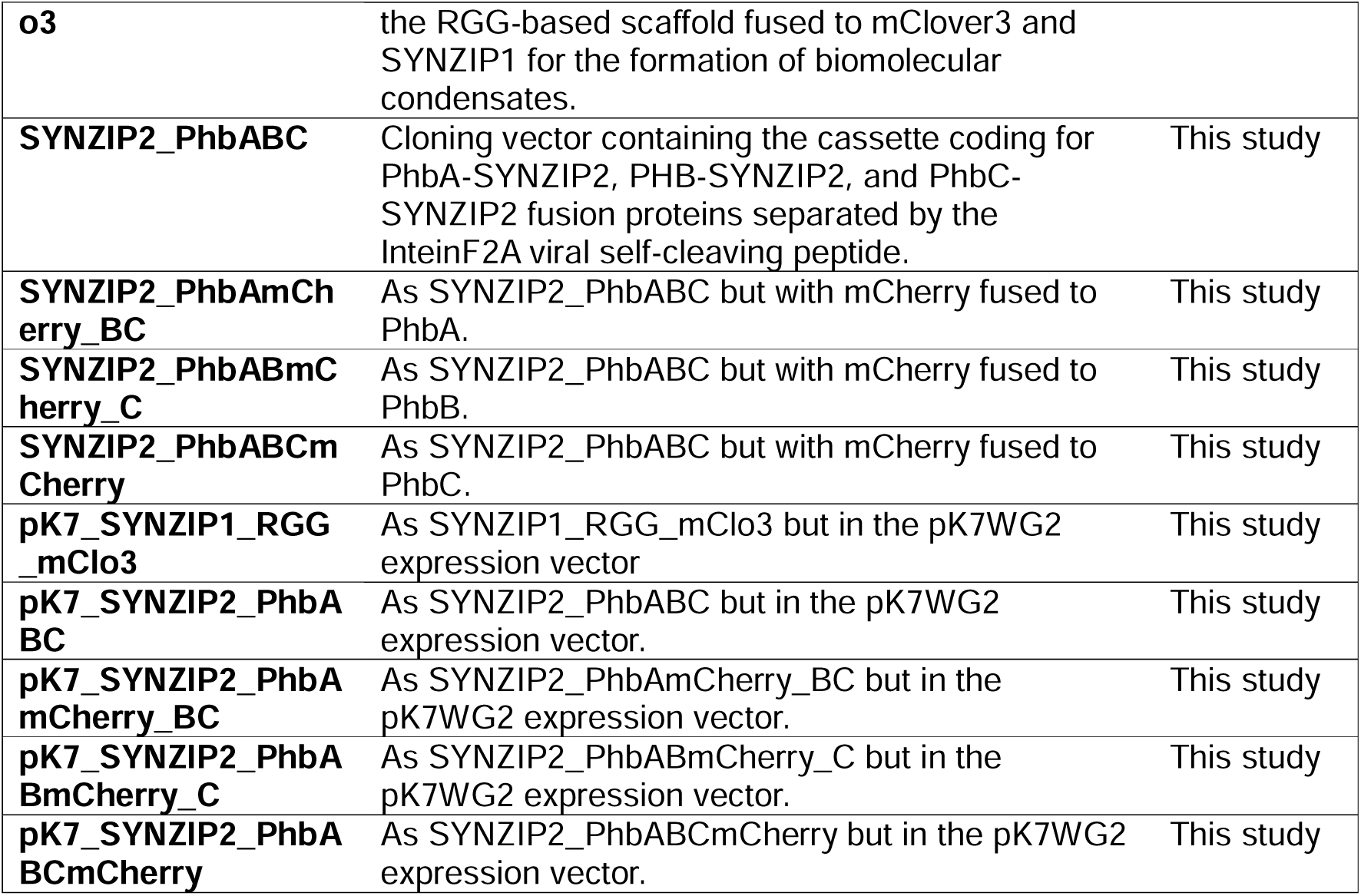
Plasmids used in this study. PhbA: β-ketothiolase, PhbB: acetoacetyl-CoA reductase, PhbC: PHB synthase.

